# *In silico* screening identifies SHPRH as a novel nucleosome acidic patch interactor

**DOI:** 10.1101/2024.06.26.600687

**Authors:** Allison M. James, Ernst W. Schmid, Johannes C. Walter, Lucas Farnung

## Abstract

Nucleosomes are the fundamental unit of eukaryotic chromatin. Diverse factors interact with nucleosomes to modulate chromatin architecture and facilitate DNA repair, replication, transcription, and other cellular processes. An important platform for chromatin binding is the H2A–H2B acidic patch. Here, we used AlphaFold-Multimer to screen over 7000 human proteins for nucleosomal acidic patch binding and identify 41 potential acidic patch binders. We determined the cryo-EM structure of one hit, SHPRH, with the nucleosome at 2.8 Å. The structure confirms the predicted acidic patch interaction, reveals that the SHPRH ATPase engages a different nucleosomal DNA location than other SF2-type ATPases, and clarifies the roles of SHPRH’s domains in nucleosome recognition. Our results illustrate the use of *in silico* screening as a high throughput method to identify specific interaction types and expands the set of potential acidic patch binding factors.

**All the screening data is freely available at:** https://predictomes.org/view/acidicpatch

---

The nucleosome, composed of an octameric histone protein core and 145-147 base pairs of DNA, represents the fundamental unit of chromatin in eukaryotic cells^1,2^. A highly conserved feature of nucleosomes is the acidic patch, an array of negatively charged residues in H2A and H2B that are situated on the nucleosome surface^3^. The acidic patch plays a critical role in coordinating intermolecular interactions that influence chromatin dynamics, and it is formed by a cluster of acidic amino acids of histone H2A (Glu56, Glu61, Glu64, Asp90, Asp91, and Glu92) and histone H2B (Glu105, Glu113)^2^ (Fig. 1a-b). Together, these residues form two depressions (1 and 2) on the surface of the H2A–H2B dimer that are separated by a small ridge^3^. Proteins that bind the acidic patch typically engage the H2A–H2B acidic patch using an arginine residue that inserts into depression 1 of the acidic patch (“arginine anchor nucleosome binding motif”)^4^. Arginine anchors often exist in a disordered loop region but can exist within a variety of structural motifs (helical and beta sheet regions), making it difficult to predict arginine anchor motifs.

**Figure 1.**
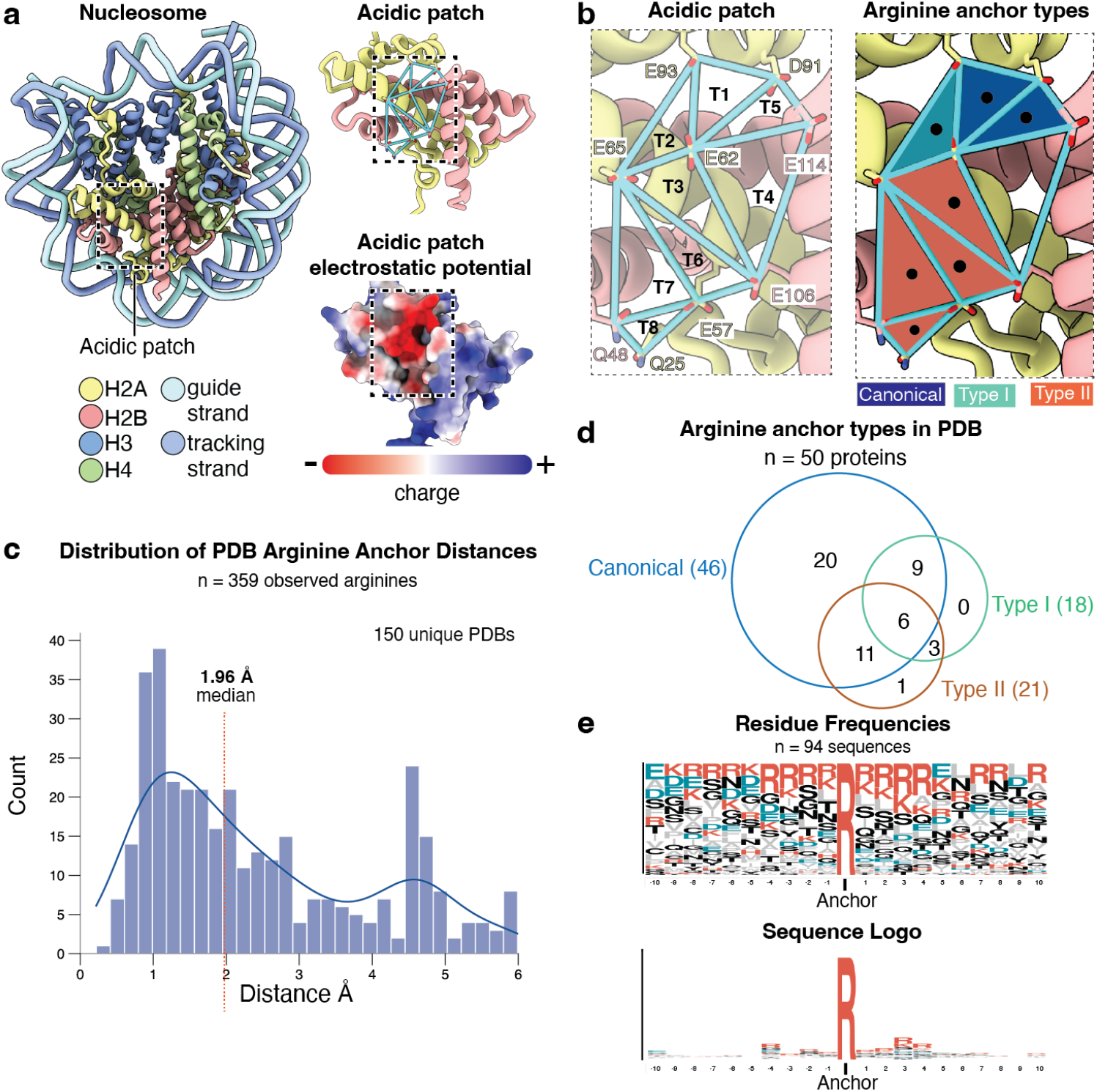
A strategy to find and evaluate acidic patch interactions. **(a)** The acidic patch is a region on the surface of the H2A–H2B histone dimer defined by a cluster of highly negatively charged residues. **(b)** A schematic showing how the acidic patch was represented using a series of 8 triangles. Triangle vertices were defined by terminal carbons on the indicated residues. **(c)** Applying the triangle-based analysis to PDB structures reveals that arginine anchors are positioned at a median of 1.96 Å away from the centers of one of the acidic patch triangles. **(d)** Based on the triangles, PDB anchors were sorted into 3 distinct binding modes (canonical, type 1, and type 2). **(e)** Sequence logo analysis of the residues around the anchoring residue reveals no distinct anchor motif with only minor enrichment for other positively charged residues (arginines and lysines).

Because the acidic patch is an essential interaction platform for many chromatin binding proteins, previous studies have identified numerous acidic patch interactors using techniques such as affinity pull-down and mass spectrometry^5,6^. However, these methods have limitations, including the inability to identify precise binding residues and difficulty detecting weak or transient interactions. For example, the ubiquitin E3 ligase RNF168, a known acidic patch binder^7^, was not identified in mass spectrometry screens.

To overcome these limitations and expand the known interactome of the acidic patch, we developed an *in silico* screening approach using AlphaFold-Multimer predictions^**8**^. This approach allowed us to examine potential interactions between the H2A-H2B dimer and over 7000 nuclear human proteins. Our screen identified over 40 novel, highconfidence acidic patch interactor candidates. To validate our approach, we focused on one of these candidates, SHPRH (SNF2, histone-linker, PHD and RING finger domaincontaining helicase), an E3 ubiquitin-protein ligase. We obtained a cryo-EM structure of SHPRH bound to a nucleosomal substrate at 2.8 Å resolution, confirming the predicted acidic patch engagement and revealing new insights into SHPRH’s domain architecture and nucleosome recognition mechanism. In sum, our *in silico* H2A–H2B screen identifies a broad range of acidic patch interactors and opens new avenues for hypothesis generation and targeted experimental validation in the chromatin field. All the screening data is freely available at https://predictomes.org/view/acidicpatch.

## Results

### In silico protein interaction screening reveals the predictome of nuclear human H2A–H2B acidic patch binders

To systematically identify acidic patch binders, we screened 7,608 nuclear *H. sapiens* proteins using AlphaFold-Multimer^9^ (Methods). Each of the 7,608 proteins was “folded” with an *H. sapiens* H2A–H2B dimer in ColabFold, and each trimeric protein combination was predicted using 3 of 5 AlphaFold-Multimer models, yielding a total of 22,824 predicted structures (Methods). Although several analysis methods have been developed to rank predictions from large scale AlphaFold multimer screens, such as pDOCKQ2^10^ and SPOC score^11^, these methods are not suitable for identifying specific types of interactions such as those that occur during acidic patch binding. We therefore created a novel analysis pipeline specifically designed to examine residue interactions required for acidic patch binding.

We developed a geometric analysis workflow in which we represented the acidic patch surface of the H2A–H2B dimer as a series of 8 inter-connected triangles (Fig. 1b; Methods). We searched the predicted structures for any arginine residue whose terminal carbon is less than 6 Å from the center of any of the triangles (the maximum distance at which electrostatic interactions are expected to impact binding). To determine if this procedure can retrieve known acidic patch binding arginine anchors, we analyzed all H2A/H2B containing structures across all species from the PDB (queried on April 1, 2024) (Methods). This yielded 150 different PDB structures containing 359 observations of arginine anchors positioned a median of 1.96 Å away from the acidic patch triangles (Fig. 1c). For a more granular classification, we categorized arginines as canonical, type I, or type II, based on the closest triangle in the acidic patch with which it interacts (Fig. 1b). Among the 50 unique proteins (excluding unassigned polypeptide entities such as nanobodies) found across the 150 PDB structures, >90% utilized a canonical anchor, suggesting that this is the dominant binding mode by which proteins engage with the acidic patch (Fig. 1d). We next asked whether the sequence context around these arginine residues could be used to identify an acidic patch anchoring motif. An analysis of 94 unique sequences spanning 10 amino acids before and after the anchoring arginine did not reveal such a motif, although there was a slight enrichment for adjacent positively charged residues (lysine and arginine residues) (Fig. 1e). These results suggest that finding novel acidic patch interactors is likely not possible by searching for a sequence motif and requires a more sophisticated approach such as the *in silico* structural interaction screen followed by the geometric analysis we devised.

Having validated our geometric analysis on the known acidic patch interactions, we next analyzed our *in silico* predictions using the same geometric analysis pipeline. This revealed 6,280 unique arginine residues in 3,364 nuclear proteins that were located within 6 Å from the center of any triangle (∼44% of all proteins screened) (Fig. 2a-b). We then assessed AlphaFold-Multimer’s confidence in the placement of these arginine residues by examining the predicted Alignment Error (PAE) output. This metric spans 0 Å (best) to ∼30 Å (worst) and quantifies the model’s confidence in the relative positions of two residues. We extracted the average PAE of putative arginine anchors relative to the residues in the 3 vertices of the nearest triangle. Plotting this metric as a function of putative anchor distance, we observed a dense core distribution centered around a mean PAE of 15 Å that likely represents randomly placed, non-interactors (Fig. 2c). Most AlphaFoldMultimer predictions of known human PDB anchors have PAE values lower than those in the core distribution. Given this, we retained candidate anchors that lie significantly outside the baseline distribution (Fig. 2d; Methods) and have mean PAEs < 15 Å (Fig. 2d).

**Figure 2.**
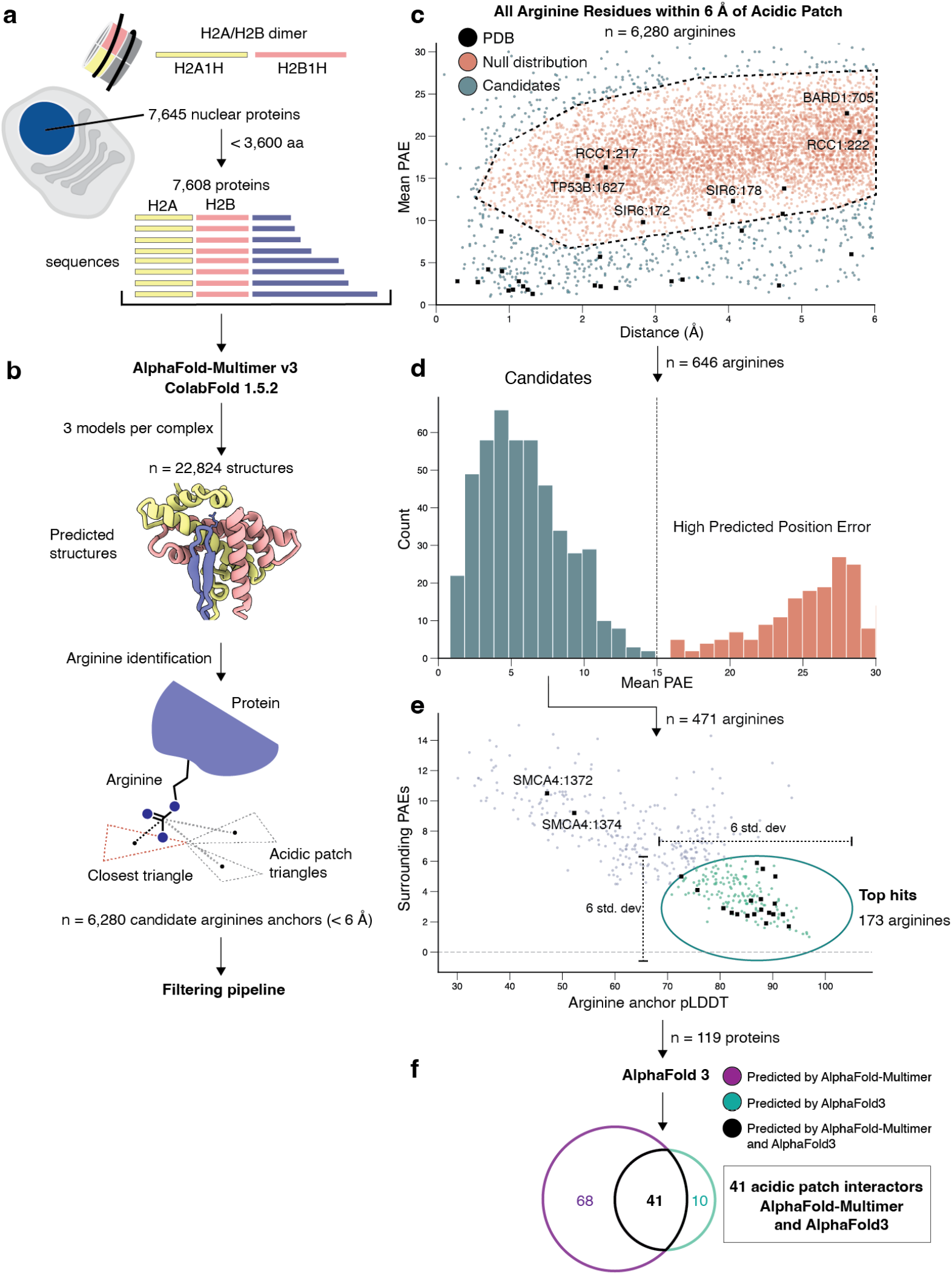
A high throughput AlphaFold-Multimer based screen to identify novel acidic patch binders. **(a)** To select candidate proteins for screening, we used UniProt to retrieve sequences for proteins associated with a nuclear localization annotation. These were filtered by length to remove any where the total sequence when concatenated to the H2A/H2B sequence exceeded the GPU memory-imposed limit of 3,600 amino acid. **(b)** Concatenated sequences were run via AlphaFold-Multimer/ColabFold and the resulting structures were analyzed to extract distance and confidence metrics associated with the predictions. **(c)** A scatter plot of all arginine anchors. A gaussian kernel was fit to the data and used to identify a core null (non-interaction) distribution. Candidate arginines (outliers from this distribution) were extracted for downstream analysis. **(d)** Examining the mean PAE associated with outlier anchors shows a bi-modal distribution. The lower error (more confident) subset was then extracted. **(e)** To further enrich for the most confident hits, we plotted the anchor residue pLDDT vs the mean PAE of anchor adjacent residues to the residues in the nearest acidic patch triangle. Superimposing the values obtained when predicting groundtruth anchors form the PDB reveals a cluster in the lower right. Anchors were isolated by defining an elliptical region centered at the PDB medians and with major and minor axes defined by the standard deviation of these two metrics. **(f)** The 119 most confident interactors/proteins were run against a full nucleosome core particle via the AlphaFold 3 server. The results were analyzed to determine if the previously identified arginines were once again predicted to interact with key acidic patch residues.

As a final filter, we determined the predicted Local Distance Difference Test (pLDDT) value of each identified arginine anchor. This confidence metric, in contrast to PAE, measures local positioning confidence of a residue relative to its immediate spatial neighbors. Graphing the pLDDT of the anchor versus its mean PAE (as defined above) reveals a cluster of anchors that is highly enriched for known PDB arginine anchors (Fig. 2e). We selected anchor residues whose pLDDT and surrounding mean PAE were within 3 standard deviations in either direction of the corresponding median values for known true positives from the PDB (Fig. 2e).

After applying the above criteria, 173 arginine anchors from 119 unique proteins remained, corresponding to an overall protein hit rate of ∼1.6%. A gene ontology (GO) analysis^12^ for molecular function revealed a significant enrichment for terms relating to DNA-binding proteins, which often use arginine residues to bind the negatively charged DNA backbone. We therefore used the newly released AlphaFold 3^13^, which can model protein-nucleic acid complexes, to predict the interaction of all 119 hits with a nucleosome core particle. We looked for instances where the arginine residues were still predicted to interact with any residues from the acidic patch (pLDDT >= 30, min PAE < 15 Å and closest inter-atom distance < 5 Å). This analysis revealed that of the s119 candidates, 41 (34 %) (Table 2) were predicted to have the same arginine anchor(s) interacting with the acidic patch (Fig. 2f). In other cases, such as with the transcription factor SPIC, the previous “arginine anchor” residues were instead predicted to interact with nucleosomal DNA (Extended Data Fig. 1d-e). Subsequent GO analysis of this new set revealed a complete absence of DNA binding terms, suggesting that our final AlphaFold 3 based filtering method removes artifactual false positives that occur when DNA is not supplied during the prediction (such as in AlphaFold-Multimer/ColabFold) (Extended Data Fig. 1a).

A comparison of the final set of 41 proteins identified as hits with the PDB set reveals that our approach captures 10 of the 25 known human anchor proteins from the PDB (Fig. 3a). GO term enrichment analysis of the hits reveals that chromatin/histone related terms are among the 14 significantly over-represented annotations identified at a False Discovery Rate (FDR) of less than 0.05 (Fig. 3b). We examined the residues adjacent to the arginine anchors identified across all the hits but again saw no evidence for a strongly defined sequence motif in these extracted regions (Extended Data Fig. 1b). We next asked how the results of our screen would differ when we fold each nuclear protein with either H2A or H2B, mirroring a more typical binary interaction screen. We folded the 41 final protein hits individually with either H2A or H2B and analyzed the results for arginine anchors. Of the 41 proteins examined, only 19 (46%) were classified as having acidic patch like interactions via this method, highlighting the importance of using a dimeric bait (H2A and H2B) to capture the maximum number of partners.

**Figure 3.**
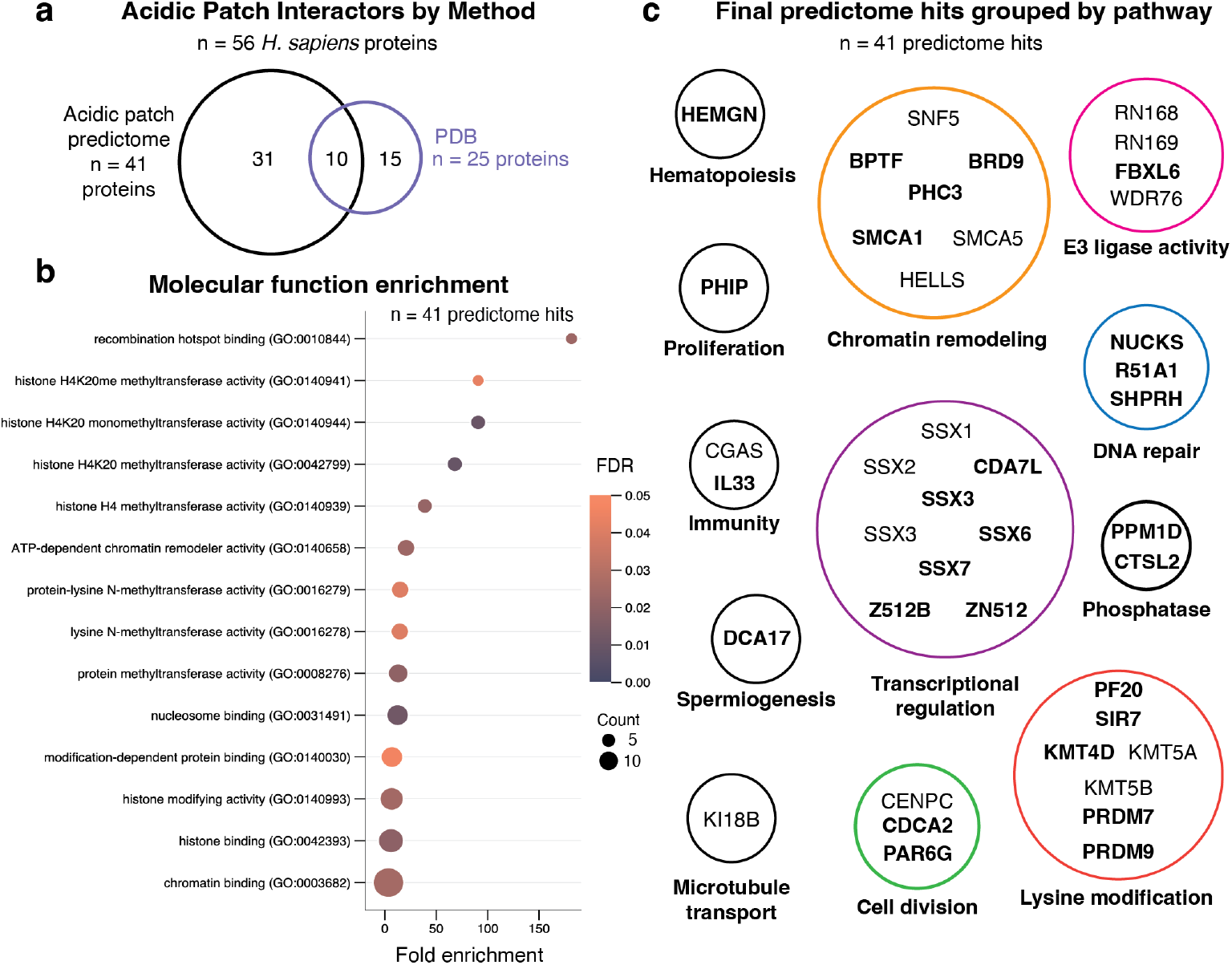
Characterization of hits identified via the predictome screen. **(a)** Venn diagram comparing the *in silico* predictome hits to those previously structurally characterized unique *H. sapiens* PDB entries. **(b)** A GO enrichment analysis for molecular function relative to the background distribution associated with the proteins run in the screen reveals terms related to nucleosome remodeling and histone modification. **(c)** Overview of all 41 proteins identified as acidic patch binders via the *in silico* predictome screen grouped by biological pathway. Novel hits with a predicted acidic patch interaction that has not been characterized structurally previously or by previous mass spectrometry screens are shown in bold.

### AlphaFold-Multimer benchmarking for acidic patch binding predictions

To assess the success rate of our screen, we compared AlphaFold-Multimer predictions of acidic patch binders from our screen with the 12 experimentally resolved structures of human acidic patch binders that were deposited in the PDB after the AlphaFold-Multimer training cutoff date. Our screen captured 6 out of these 12 proteins, and this number was reduced to 4 after our AlphaFold 3 pruning step. Thus, our *in silico* screen identified ∼33% (4/12) of the post-training human interactors, likely due to our stringent filtering criteria. Indeed, the majority of PDB-verified hits were excluded by the confidence-based filtering rather than the initial distance-based arginine search.

For the 4 known human PDB interactors not identified via the initial arginine search (BAP1, DOT1L, RING2, ZNHI1), we considered the possibility that they were falsely excluded because our AlphaFold-Multimer predictions were not relaxed and therefore the arginine side chains were not correctly positioned relative to the acidic patch. To test this idea, we performed AMBER relaxation, which settles predicted structures into more energetically favorable conformations. After re-analyzing the relaxed structures, we identified only one new potential anchor (Arg708) in the BAP1 protein^14^, and only in one of three predicted models. This anchor, however, had a pLDDT score of 20, indicating a very low confidence prediction. This result suggests that although computationally expensive structural relaxation may offer some benefits, it would not substantially alter our overall findings.

Upon closer examination of the PDB structures that were not hits in our screen, we noticed that several false negatives contained nucleosomes with posttranslational modifications. For example, BAP1 contained H2AK119Ub nucleosomes^15^ and SIRT6 contained H3K9ac marks^16^. One limitation of our approach is that we only provide unmodified H2A and H2B proteins and may miss interactions that are at least partially dependent on posttranslational histone modifications. Similarly, we may miss interactions that are mediated by additional subunits of multi-subunit complexes.

### Comparison with an available mass spectrometric acidic patch screen

To further validate our predictions, we compared our list of 41 interactors with a previous mass spectrometry screen for H2A–H2B acidic patch binders^5^. We found that 4 of our AlphaFold interactors (10%) were also identified as hits via a mass spectrometry screen^5^ (Extended Data Fig. 1c). We also examined the overlap of the PDB set of human acidic patch binders (as defined above) with the hits found via AlphaFold-Multimer or the published mass spectrometry pulldown (Table S4). Notably, our AlphaFold-Multimer screen identified 10 (40%) of all known human PDB acidic patch interactors, nearly twice as many as the mass spectrometry-based approach^5^.

### Validation of available structural data and identification of novel hits

We found excellent agreement between our predictions and PDB structures and identified a total of 41 H2A–H2B acidic patch interactor candidates. For example, we model the arginine anchors of the E3 ligase RNF168 with near-perfect agreement^7^ (Fig. 4a). Moreover, we observe excellent agreement with experimental data for the synovial sarcoma X (SSX) breakpoint protein SSX1, which was also not included in the AlphaFold-Multimer training set^17^ (Fig. 4b). Four of our hits (HELLS, WDR76, KIF18B, and SMARCA5) are identified as acidic patch interactors by mass spectrometry, and do not yet have an experimentally determined structure characterizing the acidic patch interaction. For example, we predict the arginine anchor interaction for the chromatin remodeler SMARCA5 (SMCA5/SNF2h). A SMARCA5-nucleosome complex structure was determined, but an acidic patch interaction was not resolved^18^. Photo-crosslinking and mutational experiments identified a conserved acidic patch binding motif that is important for SMARCA5’s remodeling activity *in vitro*^19^. Through our screen, we identify the same acidic patch binding motif and provide a structural model for the biochemically identified acidic patch interaction (Fig. 4c).

**Figure 4.**
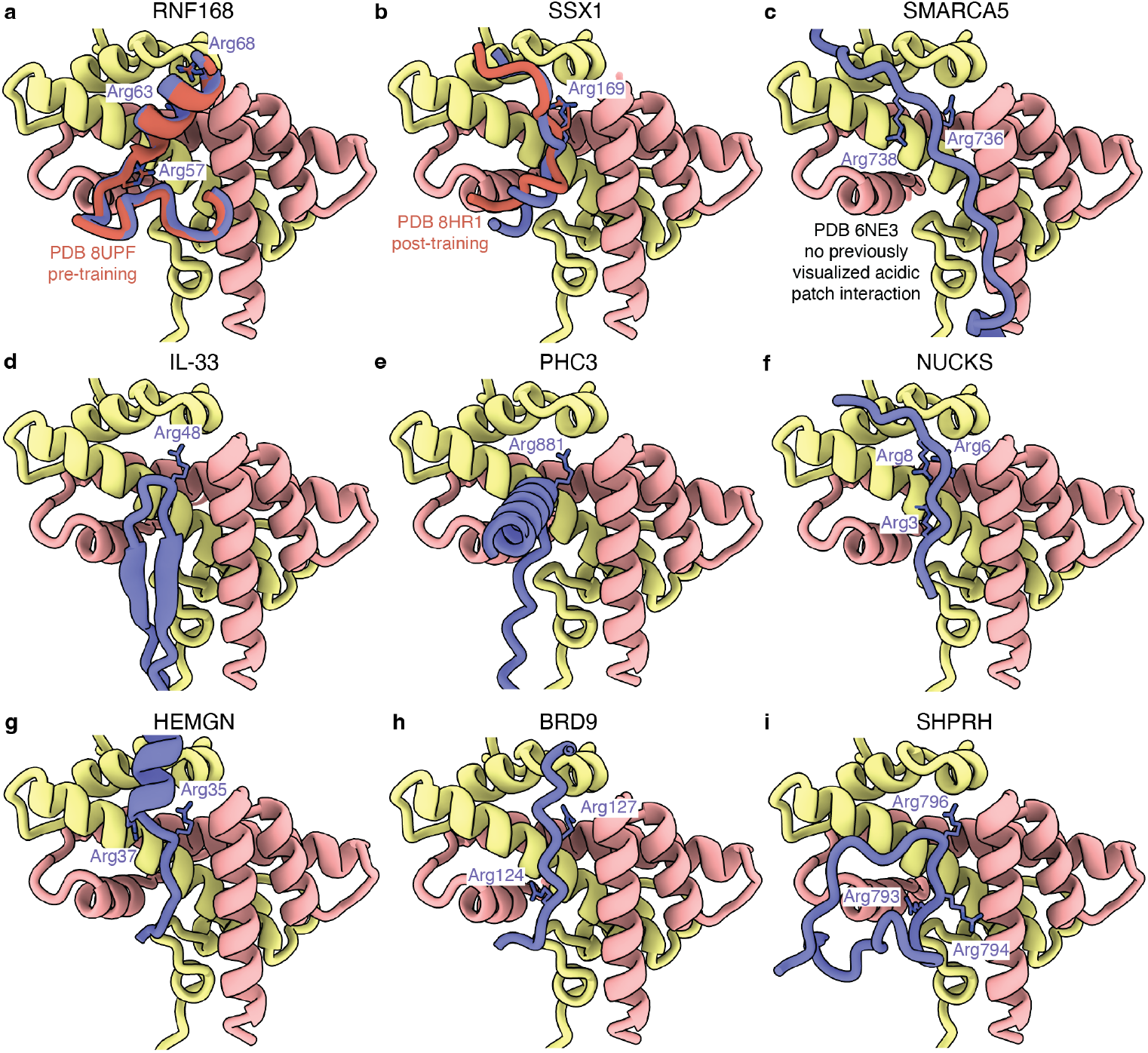
Predictome screen identifies a broad range of acidic patch interactors. **(a)** RNF168-H2A-H2B prediction. RNF168 is shown in purple, PDB 8UPF is aligned on the H2A-H2B dimer and RNF168 is overlayed in orange, H2A is shown in yellow, H2B is shown in red. Arginine anchor residues are labeled in purple. Color scheme is used throughout. **(b)** SSX1-H2A-H2B prediction. PDB 8HR1 is aligned on the H2A–H2B dimer and SSX1 is overlayed in orange (PDB 8HR1). **(c)** SMARCA5-H2A-H2B prediction. **(d)** IL-33-H2A-H2B prediction. **(e)** PHC3-H2A-H2B prediction. **(f)** NUCKS-H2A-H2B prediction. **(g)** HEMGN-H2A-H2B prediction. **(h)** BRD9-H2A-H2B prediction. **(i)** SHPRH-H2A-H2B prediction.

Finally, we identify 27 acidic patch interactors that have neither been identified by mass spectrometry nor structurally characterized. Here, we describe six of these interactors. One of our top hits is IL-33, a cytokine known to associate with chromatin^20^, and IL-33 shares sequence homology to the LANA peptide, a known acidic patch binder^21^. Our prediction is supported by previous experimental evidence showing that mutation of the identified canonical arginine anchor (Arg48) disrupts IL-33’s chromatin association^22,23^ (Fig. 4d). Another hit, PHC3, participates in a PRC1-like complex, which has a role in heterochromatin maintenance throughout development^24^ (Fig. 4e). The DNA repair factor NUCKS, which promotes homologous recombination^25^ is predicted to engage the acidic patch using all three types of arginine anchors (Fig. 4f). The hit HEMGN regulates differentiation and proliferation of hematopoietic stem cells^26^ (Fig. 4g). BRD9, a BAF complex subunit, recognizes acylated histones and plays important roles in chromatin remodeling and transcriptional regulation^27,28^ (Fig. 4h). Finally, the DNA repair factor and SF2-type ATPase protein SHPRH is a highconfidence hit and is predicted to utilize all three types of arginine anchors, suggesting a high affinity for the H2A– H2B dimer (Fig. 4i).

### Nucleosome-SHPRH complex

To validate our *in silico* predictions for SHPRH (Fig. 4i), we structurally characterized a SHPRH-nucleosome complex using single-particle cryo-EM. SHPRH has multiple domains including a SF2-type ATPase motor, HIRAN, H15, PHD, and RING domain (Fig. 4a) and has previously been shown to bind nucleosomal substrates^29^. Its mode of interaction, however, remains unclear. The presence of a SF2-type ATPase motor suggested that SHPRH could engage nucleosomes like other SF2-type ATPase motors by binding nucleosomal DNA at similar positions as found in chromatin remodelers (SHL2, 6), although we do not observe nucleosome sliding activity or ATP hydrolysis by SHPRH (Extended Data Fig. 2a-b). Our predictome screen identified that SHPRH can engage the nucleosome via an insertion in its ATPase lobe 1 where a flexible loop inserts into the H2A– H2B acidic patch using a canonical arginine anchor (Arg796) (Fig. 4i).

We recombinantly expressed and purified *H. sapiens* SHPRH and reconstituted a nucleosomal substrate with a Widom 601 sequence flanked by 60 base pairs of extranucleosomal DNA on one side. We subsequently formed the SHPRH-nucleosome complex in the presence of the ATP transition state analogue ADP ·BeF_3_, purified the complex by size exclusion chromatography, and prepared the complex for single-particle cryo-EM (Extended Data Fig. 2c-g). We collected a dataset of 16,168 micrographs with 4,009,252 particles (Extended Data Fig. 3a). Cryo-EM data analysis revealed a complex of SHPRH bound to the nucleosome from 110,867 particles at an overall resolution of 2.8 Å (FSC 0.143 criterion) (Extended Data Fig. 3b-c, Extended Data Fig. 4). We were successful in modelling all domains of SHPRH except for the RING and H15 domains (Movie S1).

### Architecture of the SHPRH-nucleosome complex

In our cryo-EM structure, SHPRH engages the nucleosome with several domains and binds the acidic patch as predicted by our screen (Fig. 5a-b). First, the ATPase motor of SHPRH engages nucleosomal DNA at super-helical location (SHL) +5 and binds the ATP analogue ADP·BeF_3_. Second, SHPRH contains two insertions within lobe 1 of its SF2-type ATPase motor. Insertion 1 (residues 390-709) encompasses both an H1/H5-like (H15) domain and a plant homeodomain (PHD) finger. As predicted by our H2A– H2B predictome screen, insertion 2 (residues 782-806) contacts the acidic patch via an arginine anchor that is conserved from yeast to humans^3^0 (Fig. 5c-d). Third, SHPRH contains a large insertion between ATPase lobe 1 and ATPase lobe 2 that contains two large helical bundles that reach across the two DNA gyres at SHL +5 and -2 and contact the opposite site of the nucleosomal disc. The helical bundle extends across H3 and H4 towards the dyad before reaching back across the nucleosomal disc to the DNA gyres and engages SHL +5 via ATPase lobe 2.

**Figure 5.**
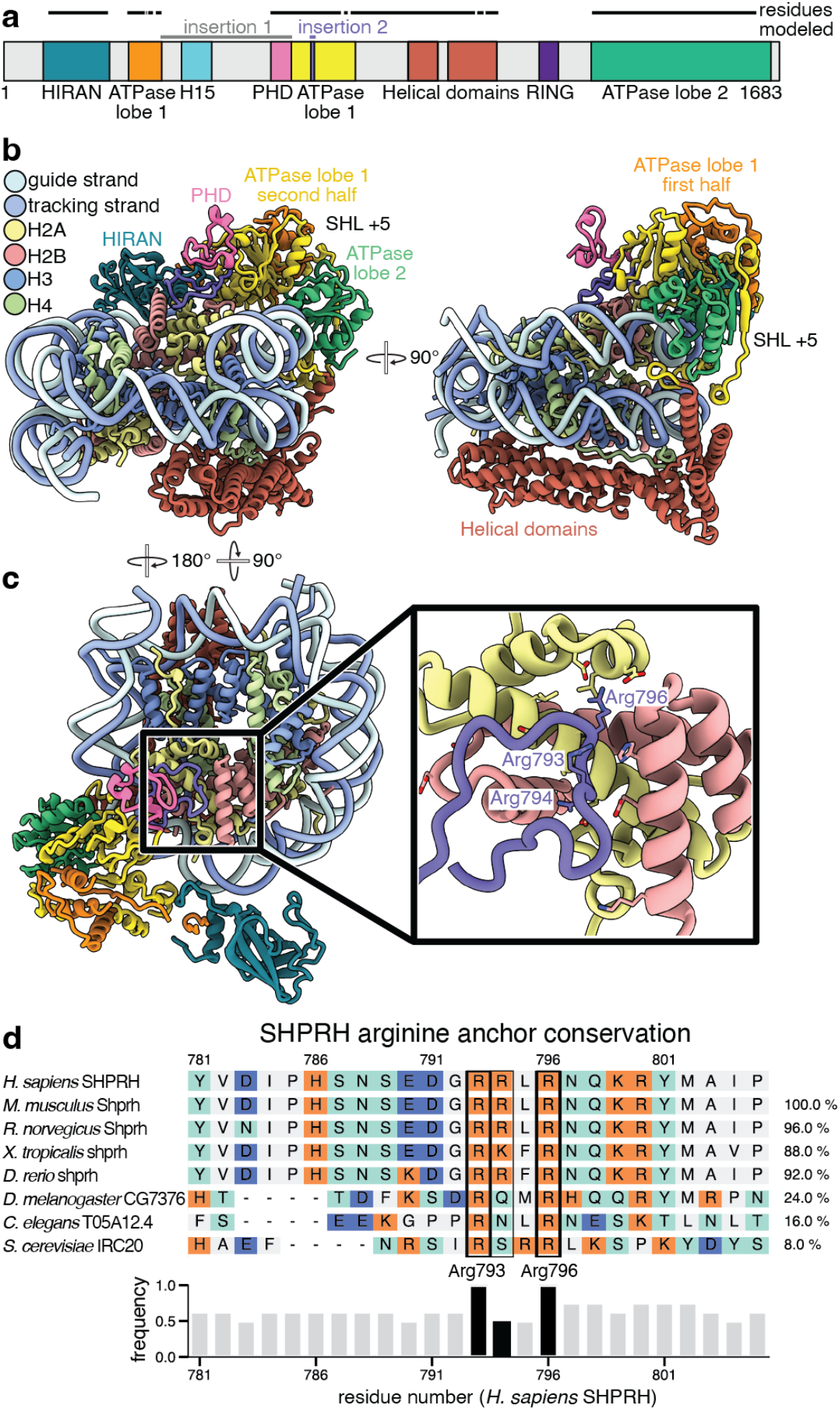
SHPRH-nucleosome structure. **(a)** Domain architecture of *H. sapiens* SHPRH. Colors used throughout. Modelled residues are indicated. **(b)** Two views of the SHPRH-nucleosome complex. **(c)** SHPRH arginine anchor (Arg796) inserts into the nucleosome acidic patch. Arg796 and the surrounding arginines (Arg793 and Arg794) are shown as sticks. **(d)** Multiple sequence alignment shows conservation of the SHPRH arginine anchor across species.

We identify a previously unannotated N-terminal HIRAN domain in SHPRH (residues 98-255) (Fig. 6a). The HIRAN domain is conserved from the yeast IRC20 ortholog and shares similarity to the HIRAN domain in yeast Rad5 and human HLTF^30–33^. The HIRAN domain sits adjacent to ATPase lobe 1 and to nucleosomal DNA at SHL +4 to +4.5 (Fig. 6a). We visualize only low-resolution density for the HIRAN domain, suggesting that it is not strongly contacting the nucleosomal DNA on our substrate.

**Figure 6.**
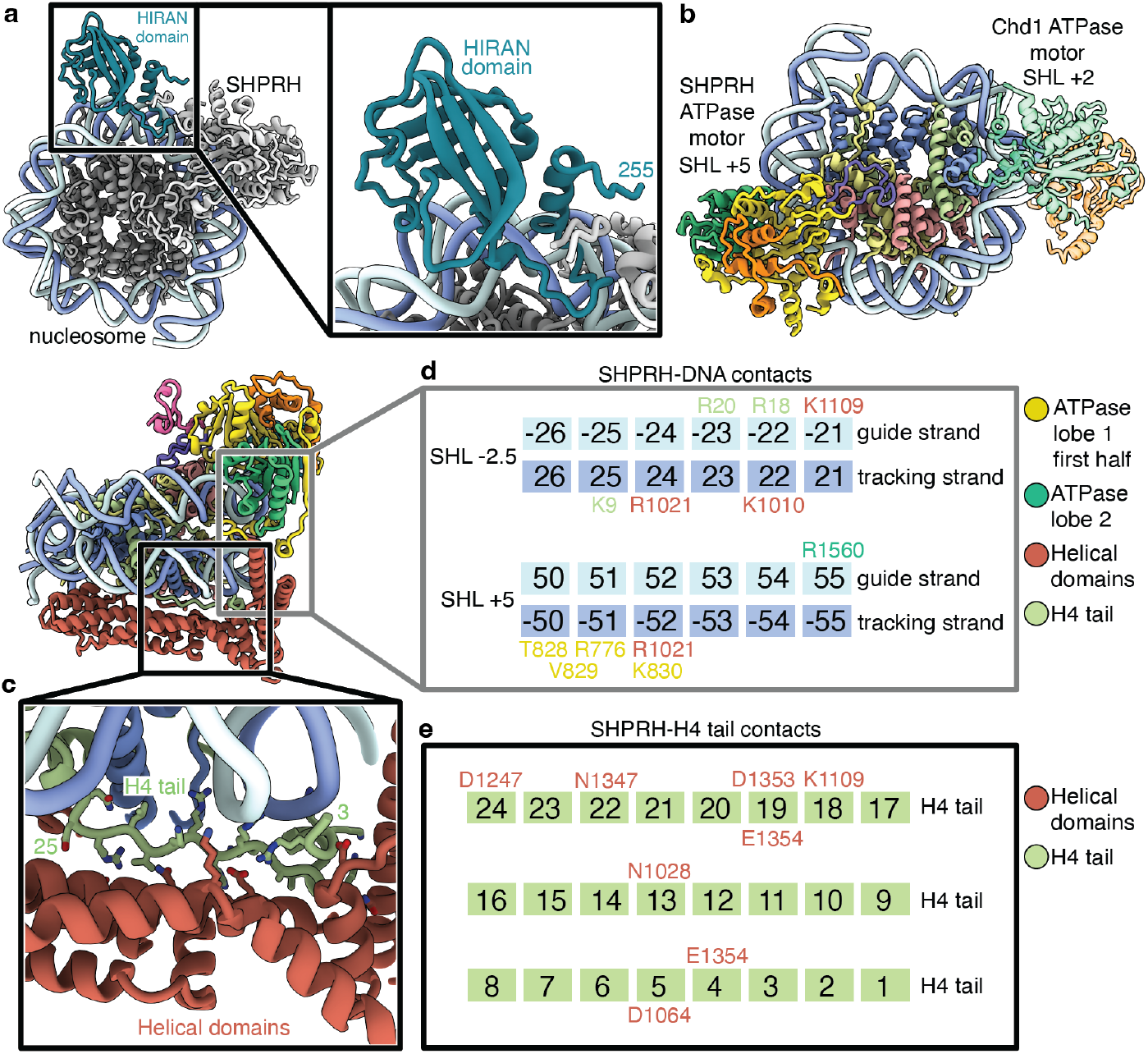
SHPRH interacts with the nucleosome via several domains. **(a)** A HIRAN domain in SHPRH sits adjacent to nucleosomal DNA at SHL +4 to +4.5. **(b)** Comparison of ATPase motor positions of SHPRH and Chd1. The ATPase motor of SHPRH contacts nucleosomal DNA at SHL +5, whereas the ATPase motor of *S. cerevisiae* Chd1 contacts nucleosomal DNA at SHL +2. **(c)** The helical domains of SHPRH contact the H4 tail (residues 3-25). The H4 tail is shown in green and side chains are shown as sticks. Heteroatom of side chains are colored. **(d)** SHPRH and the H4 tail contact nucleosomal DNA around SHL -2.5 and SHL +5. Residues contacting nucleosomal DNA are shown above or below the relevant base. **(e)** The SHPRH helical domains form contacts with the H4 tail. SHPRH residues that form hydrogen bonds with the H4 tail are indicated above or below the H4 tail residue number.

Overall, the architecture of SHPRH closely resembles the architecture predicted by our AlphaFold-Multimer screen. Alignment of the H2A–H2B dimer of our cryo-EM structure with the H2A–H2B dimer of the predicted model shows agreement in the positioning of SHPRH relative to the nucleosome with a root mean square deviation (RMSD) of 9.018 Å (Extended Data Fig. 5). This is surprising as our predictome screen did not include histones H3, H4, and nucleosomal DNA and suggests an emergent property of AlphaFold-Multimer predictions that despite the absence of structural elements still positions domains relatively accurately. Only a helical bundle of SHPRH that would clash with the H3–H4 tetramer is repositioned in our cryo-EM structure compared to the AlphaFold-Multimer based prediction. The helical bundle is correctly positioned when offered a full nucleosome core particle in AlphaFold 3, decreasing the RMSD to 3.098 Å (Extended Data Fig. 5c).

### SHPRH engagement of the H2A–H2B acidic patch mirrors predictome interactions

In agreement with our prediction that identified SHPRH residue 796 as an arginine anchor, we resolve how the arginine anchor and surrounding arginine residues (Arg793 and Arg794) engage the H2A–H2B acidic patch (Fig. 5d). Via a flexible region, insertion 2 in ATPase lobe 1 extends from nucleosomal DNA at SHL +5 towards the acidic patch and meanders back towards SHL +5. Arg796 of SHPRH acts as a canonical arginine anchor and interacts with Glu62, Asp91, and Glu93 of H2A and residues Glu106, Leu107, and His110 of H2B. Additionally, the PHD finger in insertion 1 of ATPase lobe 1 stacks on top of the acidic patch-interacting loop. Whereas many PHD fingers have been shown to recognize unmodified or methylated lysines in histone tails^34^, we are unable to visualize any PHD finger-histone tail interactions due to low resolution of the PHD domain, leaving the role of the PHD finger in SHPRH recruitment unclear. However, it is likely that the ATPase lobe 1 insertions, especially the acidic patch-interacting loop, are important to orient SHPRH on the nucleosomal substrate (Fig. 6b).

### SHPRH binds the H4 tail

ATPase lobe 1 engages one face of the nucleosome, and the SHPRH helical bundle engages the opposite nucleosomal face, and the flexible H4 tail packs between the second nucleosomal face and the helical bundle (Fig. 6c). There are extensive contacts between the H4 tail and the helical bundle of SHPRH. Additionally, SHPRH and the H4 tail show multiple contacts with nucleosomal DNA (Fig. 6d). The H4 tail inserts into the DNA major groove at SHL -1.5 via Arg20, contacts the minor groove at SHL -2 via Arg18, and again inserts into the major groove at SHL -2.5 via Lys9. The SHPRH and DNA interactions stabilize the H4 tail such that we visualize nearly the entire H4 tail (residues 3-26) with side chains, providing one of the most complete visualizations of the H4 tail to date (Fig. 6e).

## Discussion

The H2A–H2B acidic patch is a hotspot for chromatin interacting proteins such as transcription factors, chromatin remodelers, histone tail modifying enzymes, and many other enzymes important for regulating proper genome maintenance and gene expression levels^3,35^. We performed an *in silico* screen that identifies 41 proteins as high confidence nucleosome acidic patch interactors. Our screen confirms previous biochemical screens and structural data, as well as nominating 27 previously unidentified acidic patch interactors.

Our screen additionally provides an analysis platform for identifying trimeric protein interactions that would be missed in binary protein-protein AlphaFold screens. This capability is particularly valuable for understanding higherorder complexes and their interaction networks. Utilization of AlphaFold-Multimer predictions provides unique advantages over previous biochemical screens. Specifically, we identify the precise arginine anchors that interact with the acidic patch and screen the entire human nuclear proteome to identify lowly expressed or cell type specific acidic patch interactors that would be missed by other approaches. We also identify the specific subunit responsible for acidic patch interactions within multi-subunit complexes and potential transient acidic patch interactions that would be lost in pull-down approaches. Finally, the initial computational screen was accomplished in 8 days, demonstrating the efficiency and rapid turnaround of this *in silico* approach to generate new opportunities for hypothesis-driven research. Our results highlight the challenge of balancing sensitivity and specificity in computational prediction methods. While our approach effectively minimizes false positives, it may miss some true interactions due to stringent filtering criteria. We provide our unfiltered results so that others may apply their own sets of filters and encourage further exploration and analysis. Our analysis was exclusively focused on finding proteins that engage with nucleosomes via an arginine anchor insertion into the acidic patch. We expect that many other classes of nucleosomal interactions such as H2A–H2B histone chaperone interactions are contained within the dataset and encourage further exploration and analysis.

We experimentally validate one of the proteins (SHPRH) identified via the predictome screen by determining the SHPRH-nucleosome complex structure by cryo-EM. We visualize the arginine anchor insertion into the acidic patch as predicted by the screen and find excellent agreement between our experimental structural data and the predicted model. Interestingly, we find that the putative orphan chromatin remodeler SHPRH binds the nucleosome in a unique conformation that is distinct from other SF2-type ATPase containing proteins, such as Chd1 (Fig. 6b) and other chromatin remodelers^36–42^ (Extended Data Fig. 6), suggesting that at least in our observed conformation SHPRH does not perform canonical chromatin remodeling. Additionally, we visualize a previously unidentified HIRAN domain in SHPRH. Interactions between the helical domains of SHPRH and the H4 tail provide one of the most complete visualizations of the H4 tail to date (residues 3-25). Through our structural characterization of the SHPRH-nucleosome complex, we identify many nucleosome interaction surfaces and validate the predicted H2A–H2B acidic patch interaction.

In summary, the predictome database we present serves as a valuable resource for future research in the chromatin field with broad-ranging applications to mutational studies, structural characterization, and drug design across chromatin interacting protein and protein complexes.

## Acknowledgements

We thank all members of the Farnung and Walter labs for discussion. We thank The Harvard Cryo-EM Center for Structural Biology at Harvard Medical School for support with data collection. L.F. is supported by The Smith Family Awards Program for Excellence in Biomedical Research, the Damon Runyon Rachleff Innovator Award, the Rita Allen Foundation, and the NIH Director’s New Innovator Award (DP2-ES036404). E.W.S. was supported by the National Science Foundation (DGE 2140743)

## Author Contributions

L.F. cloned SHPRH. A.M.J purified protein components. A.M.J. conducted all biochemical experiments, prepared the nucleosome-SHPRH complex for cryo-EM, and processed all cryo-EM data. E.W.S. performed all PDB analysis, developed the data analysis pipeline along with the AlphaFold-Multimer screening pipeline utilized for the H2A/H2B screen. L.F. and J.C.W. designed and supervised research. L.F., E.W.S., and A.M.J. wrote the manuscript with input from all authors.

## Correspondence

should be addressed to J.C.W. and L.F.

## Materials and Methods

### Protein retrieval

Using the UniProt database^43^, we identified 7,645 *H. sapiens* proteins associated with the GO term for the cellular component nucleus (GO:0005634) on April 4^th^, 2023. We then selected the canonical isoform of each of these protein as provided by UniProt and included these sequences in our screen. To avoid GPU memory exhaustion, all proteins that together with the H2A–H2B dimer exceeded 3,600 amino acid residues were excluded. The remaining 7,608 unique protein sequences were folded against an *H. sapiens* H2A–H2B dimer using ColabFold. H2A type 1H (Q96KK5) was used as the representative human H2A sequence and H2B type 1H (Q93079) was used as the representative human H2B sequence. The exact sequences run are found in (Table S2).

### Acidic patch geometric (triangle) analysis

A script was written in Python to parse PDB or CIF files and construct one or more three dimensional triangles with vertex coordinates extracted from specific atoms in the structures. Triangles are specified by supplying the script with one or more protein sequence strings that indicate which residues in the structure will supply atoms to use as triangle vertices. These vertices are found by aligning the supplied sequence strings to the amino acid sequences in the structure file and choosing the alignment with the most identities with a minimum required identity of 60%. This allows the script to handle cases where there are no exact sequence matches, as is the case when searching for the acidic patch regions in histone variants across the PDB. Once vertices are found and triangles are defined, the script then searches for any arginine (not in the same chain as any of the vertices) with a terminal carbon that is closest to the center of the triangle. If this distance is less than 6 Å, this arginine is identified as an anchor. If the structure being analyzed is an AlphaFold-Multimer prediction, the script will additionally extract pLDDT information as well as PAE information from the PAE JSON file that corresponds to the structure file undergoing analysis. The following 8 strings were used to define the 8 triangles used during acidic patch analysis of PDB and AFM structures : (T1: “AIRND*EE*LNK; YLTAE*ILELAG”, T2: “AIRND*EELNK; YLTAE*ILE*LAG”, T3: “YLTAE*ILE*LAG; HAVSE*GTKAV”, T4: “YLTAE*ILELAG; PGE*LAKHAVSE*GTK”, T5: “YLTAE*ILELAG; AIRN-DEE*LNK;PGE*LAKHAV”, T6: “AAVLE*YLTAEILE*LAG; HAVSE*GTKAV”, T7: “AAVLE*YLTAEILE*LAG; KVLKQ*VHPDT”, T8: “SSRAGLQ*FP; AAVLE*YLTAE; KVLKQ*VHPDT”).

### PDB arginine anchor retrieval

First all H2A/H2B containing structures in the PDB were isolated by performing a sequence search via the PDB API for any structures containing protein chains with sequences at least 70% identical to N-APVYM AAVLE YLTAE ILELA GNAAR DNKKT-C (H2A fragment) and N-TSREI QTAVR LLLPG ELAKH AVSEG TKAVT-C (H2B fragment) with an E-value < 0.1. CIF structure files corresponding to the final structure model for these entries were then downloaded and analyzed for arginine anchor based acidic patch binding via the triangle analysis script. We restricted our search to arginine residues that are flanked on both sides by at least 3 amino acids. Using our triangle analysis, we identified 47 solved unique acidic patch anchors spanning 25 human proteins, which we refer to as “the PDB set”.

Structures were characterized as either pre or post AlphaFold-Multimer v3 training if their release date was either before (pre-training) or after 2021-09-30 (post-training), respectively. Of these 25, 12 proteins were found post-training, i.e., they were found in structures deposited in the PDB after training (October 2021). Newly deposited structures with an acidic patch interaction that were already observed prior to October 2021 were excluded from our post-training set. Based on the structures, we identified the arginine anchor that binds the depression 1 of the acidic patch for each of the structures and categorized if a structure of the same factor bound to the nucleosome already existed prior to the training cutoff date. All analysis results are recorded in (Table S1).

### Sequence motif analysis and figure generation

Sequences flanking identified arginine anchors were extracted from structure files (PDB or CIF), deduplicated and then submitted to the WebLogo online server^44^ for visualization and analysis.

### AlphaFold-Multimer screening

AlphaFold was run via a customized version of ColabFold v1.5^45^ running locally on a Linux server equipped with 8 A100 GPUs rented from Lambda Labs cloud services. Predictions were generated using AlphaFold-Multimer version 3 weights (models 1, 2, 4) with monomeric templates enabled, dropout disabled, 1 ensemble, 3 recycles, and no AMBER relaxation. Unpaired + paired MSAs were supplied to AlphaFold-Multimer via the ColabFold MSA generation pipeline which routes requests to an external secondary server that runs the MMSeq2 sequence alignment algorithm^46^. In every prediction chain A and chain B are H2A1H and H2B1H respectively while chain C is variable and represents the protein run against the H2A/H2B dimer. Protein sequences used are recorded in (Table S2).

### AlphaFold-Multimer confidence-based hit pruning

Hits initially identified via the multimer based analysis pipeline were then re-run using the AlphaFold 3 web server. Each of the proteins was input as a single copy as protein chain A along with a full nucleosome core particle (Xenopus *laevis* histones + 2 strands from the Widom 60 DNA sequence) constituting the remaining chains B -K. CHAINS B/C H2A: MSGRG KQGGK TRAKA KTRSS RAGLQ FPVGR VHRLL RKGNY AERVG AGAPV YLAAV LEYLT AEILE LAGNA ARDNK KTRII PRHLQ LAVRN DEELN KLLGR VTIAQ GGVLP NIQSV LLPKK TESSK SAKSK. CHAINS D/E H2B: MAKSA PAPKK GSKKA VTKTQ KKDGK KRRKT RKESY AIYVY KVLKQ VHPDT GISSK AMSIM NSFVN DVFER IAGEA SRLAH YNKRS TITSR EIQTA VRLLL PGELA KHAVS EGTKA VTKYT SAK. CHAINS F/G H3: MARTK QTARK STGGK APRKQ LATKA ARKSA PATGG VKKPH RYRPG TVALR EIRRY QKSTE LLIRK LPFQR LVREI AQDFK TDLRF QSSAV MALQE ASEAY LVALF EDTNL AAIHA KRVTI MPKDI QLARR IRGER A. CHAINS H/I: MSGRG KGGKG LGKGG AKRHR KVLRD NIQGI TKPAI RRLAR RGGVK RISGL IYEET RGVLK VFLEN VIRDA VTYTE HAKRK TVTAM DVVYA LKRQG RTLYG FGG. CHAIN J: 5’-ATC GAT GTA TAT ATC TGA CAC GTG CCT GGA GAC TAG GGA GTA ATC CCC TTG GCG GTT AAA ACG CGG GGG ACA GCG CGT ACG TGC GTT TAA GCG GTG CTA GAG CTG TCT ACG ACC AAT TGA GCG GCC TCG GCA CCG GGA TTC TGA T-3’. CHAIN K: 5’-ATC AGA ATC CCG GTG CCG AGG CCG CTC AAT TGG TCG TAG ACA GCT CTA GCA CCG CTT AAA CGC ACG TAC GCG CTG TCC CCC GCG TTT TAA CCG CCA AGG GGA TTA CTC CCT AGT CTC CAG GCA CGT GTC AGA TAT ATA CAT CGA T-3’.

### AlphaFold 3 acidic patch analysis

Acidic patch binding was assessed by first extracting all arginines with pLDDTs >= 30 and any atoms within 5 Å of any atoms from key acidic patch residues within the predicted PDB files. For this analysis, acidic patch residues were defined as residues (Q25, Q45, E57, E65, D91, E93) in chains (B/C) Xenopus *laevis* H2A and residues (E103, E111) in chains (D/E) Xenopus *laevis* H2B. These initial contacts were then filtered by removing any residue-pairs with mean inter-atom PAEs > 15 Å. All unique remaining arginines were classified as acidic patch anchor residues. These arginines were then cross-referenced with the initial list generated by the AlphaFold multimer filtering protocol. Only those anchors that were identified via both methods were selected as the final hits. Results and intermediate data are recorded in (Table S3).

### GO term enrichment analysis

GO enrichment analysis was performed by using the online enrichment tool found at https://geneontology.org/. Protein IDs were supplied in the form UniProt IDs. All enrichment scores were determined relative to the background protein set defined as the 7,608 proteins run via AlphaFold multimer during the initial acidic patch anchor screen.

### SHPRH ortholog sequence alignment generation

Ortholog sequences were fetched and aligned to the canonical human SHPRH sequence (Q149N8) using the web tool (https://www.flyrnai.org/cgi-bin/DRSC_orthologs.pl)^47^.

### Cloning & expression

*H. sapiens* SHPRH was cloned into LIC-compatible vectors from cDNA and was expressed in insect cells. The construct contains an N-terminal 6×His tag followed by a maltose binding protein (MBP) tag and a tobacco etch virus protease cleavage site. Bacmid, virus, and protein production were performed as previously described^36^. *X. laevis* histones were expressed and purified as previously described^36,48^. Hi5 cells (600 ml) grown in ESF-921 media (Expression Systems) were infected with 300 μL of V1 virus for protein expression. The cells were grown for 48–72 h at 27 °C. Cells were harvested by centrifugation (238*g*, 4 °C, 30 min) and resuspended in lysis buffer (300 mM NaCl, 20 mM Na·HEPES pH 7.4, 10% (v/v) glycerol, 1 mM DTT, 30 mM imidazole pH 8.0, 0.284 μg ml−l leupeptin, 1.37 μg ml−1 pepstatin A, 0.17 mg ml−1 PMSF, 0.33 mg ml−1 benzamidine). The cell resuspension was snap frozen and stored at −80 °C.

### Purifications

*H. sapiens* SHPRH and *S. cerevisiae* Chd1 (residues 1-1274) were purified in the same manner. Cells expressing SHPRH or Chd1 were resuspended in lysis buffer (300 mM NaCl, 20 mM Na HEPES pH 7.4, 10% (v/v) glycerol, 30 mM imidazole, 1 mM TCEP, 0.284 µg ml–1 leupeptin, 1.37 µg ml–1 pepstatin A, 0.17 mg ml–1 PMSF and 0.33 mg ml–1 benzamidine) and purification was performed at 4°C. Cells were subsequently lysed by sonication and the lysate was centrifuged and cleared by ultra-centrifugation. The supernatant containing SHPRH or Chd1 was subsequently filtered using 0.8 µm syringe filters. The filtered supernatant was applied to a HisTrap HP 5 mL (Cytiva). The column was subsequently washed with 10 CV lysis buffer, 3 CV high salt buffer (1000 mM NaCl, 20 mM Na HEPES pH 7.4, 10% (v/v) glycerol, 30 mM imidazole, 5 mM b-mercaptoethanol, 0.284 µg ml–1 leupeptin, 1.37 µg ml–1 pepstatin A, 0.17 mg ml–1 PMSF and 0.33 mg ml–1 benzamidine), and 5 CV lysis buffer. A self-packed XK column (Cytiva) with 15 mL of Amylose resin (New England Biolabs) was attached to the HisTrap column. The protein of interest was eluted from the HisTrap column using nickel elution buffer (300 mM NaCl, 20 mM Na HEPES pH 7.4, 10% (v/v) glycerol, 500 mM imidazole, 5 mM b-mercaptoethanol, 0.284 µg ml–1 leupeptin, 1.37 µg ml– 1 pepstatin A, 0.17 mg ml–1 PMSF and 0.33 mg ml–1 benzamidine). The HisTrap column was removed, and the amylose column was washed with 5 CV lysis buffer. The protein of interest was then eluted from the amylose column with 5 CV amylose elution buffer (300 mM NaCl, 20 mM Na HEPES pH 7.4, 10% (v/v) glycerol, 30 mM imidazole, 116.9 mM maltose, 5 mM b-mercaptoethanol, 0.284 µg ml–1 leupeptin, 1.37 µg ml–1 pepstatin A, 0.17 mg ml–1 PMSF and 0.33 mg ml–1 benzamidine). The elution was fractionated and analyzed using SDS-PAGE. Fractions containing SHPRH or Chd1 were pooled and applied to dialysis in dialysis buffer overnight (300 mM NaCl, 20 mM Na HEPES pH 7.4, 10% (v/v) glycerol, 30 mM imidazole, 5 mM b-mercaptoethanol, 0.284 µg ml–1 leupeptin, 1.37 µg ml– 1 pepstatin A, 0.17 mg ml–1 PMSF and 0.33 mg ml–1 benzamidine). 1.5 mg of TEV protease was added to the sample prior to dialysis to remove the N-terminal His6-MBP tag.

The dialyzed sample was applied to a HisTrap HP 5 mL, pre-equilibrated in lysis buffer. The flow-through containing SHPRH was collected and subsequently concentrated using an Amicon 100,000 MWCO centrifugal filter unit (Millipore). The concentrated sample was applied to a Superose 6 Increase 10/300 GL (Cytiva), equilibrated in gel filtration buffer (300 mM NaCl, 20 mM Na HEPES pH 7.4, 10% (v/v) glycerol, 1 mM TCEP). The elution was fractionated and analyzed by SDS-PAGE. Sample containing SHPRH or Chd1 was concentrated using Amicon 100,000 MWCO centrifugal filter unit (Millipore). All concentrated sample were subsequently aliquoted, flash-frozen, and stored at -80 °C prior to future use.

### Octamer formation and nucleosome reconstitution

*X. laevis* histones were expressed and purified as described. DNA fragments for nucleosome reconstitution were generated by PCR as described^36^. A vector containing the Widom 601 sequence was used as a template for PCR. Large-scale PCR reactions were performed with two PCR primers (60w0 construct: forward primer: 5’-CTA CAT TCC AGG CAG TGC CTC TGC CGC CGG CCT GTT ATT CCT AGT AAT CAA TCA GTG CCT ATC GAT GTA TAT ATC TGA CAC GTG CCT-3’, reverse primer: 5’-6FAM/ ATC AGA ATC CCG GTG CCG-3’; 30w30 construct: forward primer: 5’-CCT GTT ATT CCT AGT AAT CAA TCA GTG CCT ATC GAT GTA TAT ATC TGA CAC GTG CCT-3’, reverse primer: 5’-6FAM/ CAA CTA AAG CTT AGA TGT GCG AAT TCC AGC CAT CAG AAT CCC GGT GCC G-3’) at a scale of 25 mL. Nucleosome core particle reconstitution was performed using the salt-gradient dialysis method^48^. Quantification of the reconstituted nucleosome was achieved by measuring absorbance at 280 nm. Molar extinction coefficients were determined for protein and nucleic acid components and were summed to yield a molar extinction coefficient for the reconstituted extended nucleosome. Nucleosomes are fluorescently labelled as indicated by the primers used to generate the nucleosomal DNA substrates.

### Sample preparation for cryo-EM

SHPRH-NCP complexes were formed by incubating 3.6 µM SHPRH, 1.7 µM NCP containing 60bp of extranucleoomal DNA on one side, and 1 mM ADP·BeF_x_ in buffer containing 50 mM NaCl, 20 mM Na•HEPES pH 7.4, 2 mM MgCl_2_, and 4% glycerol on ice for 20 mins. The sample was centrifuged for 10 mins at 21,000xg for 10 mins to remove any precipitate. The complex was purified by gel filtration using a Superose 6 Increase 3.2/300 GL column. Fraction A from the gel filtration was mildly crosslinked with 0.1% glutaraldehyde for 10 minutes at 4°C before being quenched with 2 mM lysine and 8 mM aspartate. 30 µL of the sample was then dialyzed in a 30,000 MWCO dialysis cup against 500 mL dialysis buffer (50 mM NaCl, 20 mM Na•HEPES pH 7.4, 2 mM MgCl_2_) for 3 hours at 4°C.

The SHPRH-NCP complex was then frozen on Quantifoil R2/1 on 200 Mesh Copper grids that were glow discharged for 30 s at 15 mA with 10s hold time using a Pelco Easiglow plasma discharge system. 2 μL of sample were applied on each side of the grid, incubated for 8 s, blotted with Ted Pella standard vitrobot filter paper for 5 s with blot force 8 and vitrified by plunging into liquid ethane using a Vitrobot Mark IV (FEI Company), operated at 4°C and 100 % humidity.

### Cryo-EM data collection & analysis

Grids were imaged and data was collected on a ThermoFisher Scientific Titan Krios operated at 300 keV equipped with a Gatan BioQuantum GIF and a Gatan K3 direct electron detector. Data acquisition was automated using SerialEM (v3.8.6) software at a nominal magnification of 105,000, corresponding to a pixel size of 0.83 Å in nanoprobe EFTEM mode. Movies consisting of 50 frames were collected in counted mode with 2.497 s exposure time and total exposure of 51.11 e^-^/Å^2^.

Image processing and analysis were performed with cryoSPARC (v3.3.2). Movies were aligned using patch motion correction followed by contrast transfer function (CTF) estimation in cryoSPARC. Particles were picked by blob-based automatic picking, resulting in 4,009,252 particles from 16,168 micrographs. Particles were extracted with a box size of 360^2^ pixels. All classifications and refinements were conducted in cryoSPARC. Initial 2D classification was used to select particles containing nucleosome-like density. Three ab-initio volumes revealed SHPRH-nucleosome, nucleosome alone, and junk classes (Extended Data Fig. 3). Heterogeneous refinements were used to sort SHPRH-bound nucleosomes from unbound nucleosomes and low-resolution particles. Non-uniform refinement of selected particles (110,867 particles) was performed, resulting in a 2.8 Å map of the SHPRH-nucleosome complex (map A).

DeepEMhancer map sharpening^49^ was performed to obtain a sharpened map used for model building and refinement (map B). Local resolution estimation was performed in cryoSPARC using a FSC threshold of 0.143 and local resolution visualization was performed in UCSF ChimeraX.

### Model building & refinement

ModelAngelo^50^ was used to build an initial atomic model of SHPRH and histone proteins into the cryo-EM reconstruction. DNA was docked from an available nucleosome crystal structure into the density (PDB 3LZ0) and extranucleosomal DNA was manually built in UCSF ChimeraX and COOT (version 0.9). Regions not modelled by ModelAngelo were completed manually in COOT (version 0.9) and the atomic model of the SHPRH-nucleosome complex was subsequently adjusted using ISOLDE (version 1.6) and in UCSF ChimeraX. ADP·BeF_3_ and a coordinated Mg^2+^ ion was placed into the corresponding density by aligning PDB 5O9G to ATPase lobe 1 of SHPRH in UCSF ChimeraX and docking the molecules into their corresponding density. Where the cryo-EM map resolution permitted, the model was inspected residue-by-residue using ISOLDE and modified to improve the fit to the map while maintaining favorable geometry. In places where our resolution approached 5.0 Å or worse (for example, the HIRAN and PHD domains), we limited the building process mainly to rigid body docking of AlphaFold structures and resolving clashes. The complete model was then real space refined in PHENIX^51^ with one macro-cycle of global minimization, local rotamer fitting, morphing, and ADP refinement with an overall weight of 0.5 and an ADP individual isotropic restraints weight of 0.5. Iterative rounds of manual rebuilding in ISOLDE together with real space refinement in PHENIX resulted in final models with good geometry (Table 3, Table S5).

**Table 1.** All acidic patch interacting arginines found in the PDB.

| PDB | PDB release | AFMT | Chain | Residue index | Triangle | Distance | Anchor type | UniProt ID | Protein Name | UniProt Residue Index |
| --- | --- | --- | --- | --- | --- | --- | --- | --- | --- | --- |
| 1ZLA | 2/28/06 | pre | K | 9 | T1.1 | 1.13 | C | Q9DUM3 | Q9DUM3_HHV8 | 9 |
| 3MVD | 8/25/10 | pre | K | 217 | T3.1 | 1.68 | 2 | P25171 | RCC1_DROME | 216 |
| 3MVD | 8/25/10 | pre | K | 224 | T1.1 | 0.95 | C | P25171 | RCC1_DROME | 223 |
| 3MVD | 8/25/10 | pre | L | 217 | T3.2 | 1.65 | 2 | P25171 | RCC1_DROME | 216 |
| 3MVD | 8/25/10 | pre | L | 224 | T1.2 | 1.05 | C | P25171 | RCC1_DROME | 223 |
| 3TU4 | 11/23/11 | pre | K | 28 | T6.2 | 1.66 | 2 | P06701 | SIR3_YEAST | 28 |
| 3TU4 | 11/23/11 | pre | K | 29 | T1.2 | 1.02 | C | P06701 | SIR3_YEAST | 29 |
| 3TU4 | 11/23/11 | pre | K | 32 | T5.2 | 4.47 | C | P06701 | SIR3_YEAST | 32 |
| 3TU4 | 11/23/11 | pre | L | 28 | T6.1 | 2.31 | 2 | P06701 | SIR3_YEAST | 28 |
| 3TU4 | 11/23/11 | pre | L | 29 | T1.1 | 0.98 | C | P06701 | SIR3_YEAST | 29 |
| 3TU4 | 11/23/11 | pre | L | 32 | T5.1 | 4.59 | C | P06701 | SIR3_YEAST | 32 |
| 4JJN | 5/15/13 | pre | I | 30 | T3.2 | 3.82 | 2 | P06701 | SIR3_YEAST | 30 |
| 4KUD | 8/7/13 | pre | K | 27 | T6.2 | 1.45 | 2 | P06701 | SIR3_YEAST | 28 |
| 4KUD | 8/7/13 | pre | K | 31 | T1.2 | 1.03 | C | P06701 | SIR3_YEAST | 32 |
| 4KUD | 8/7/13 | pre | L | 27 | T6.1 | 1.73 | 2 | P06701 | SIR3_YEAST | 28 |
| 4KUD | 8/7/13 | pre | L | 31 | T1.1 | 1.14 | C | P06701 | SIR3_YEAST | 32 |
| 4R8P | 11/5/14 | pre | L | 98 | T1.2 | 0.71 | C | Q99496 | RING2_HUMAN | 98 |
| 4R8P | 11/5/14 | pre | N | 98 | T1.1 | 1.02 | C | Q99496 | RING2_HUMAN | 98 |
| 4X23 | 12/10/14 | pre | K | 8 | T1.1 | 1.59 | C | Q66LH7 | Q66LH7_RAT | 717 |
| 4X23 | 12/10/14 | pre | K | 10 | T2.1 | 2.24 | 1 | Q66LH7 | Q66LH7_RAT | 719 |
| 5E5A | 2/3/16 | pre | K | 11 | T1.1 | 1.15 | C | P03169 | VIE1_HCMVT | 486 |
| 4ZUX | 2/24/16 | pre | W | 91 | T3.2 | 1.27 | 2 | Q03067 | SGF11_YEAST | 91 |
| 4ZUX | 2/24/16 | pre | BA | 91 | T3.1 | 0.82 | 2 | Q03067 | SGF11_YEAST | 91 |
| 4ZUX | 2/24/16 | pre | W | 84 | T6.2 | 1.11 | 2 | Q03067 | SGF11_YEAST | 84 |
| 4ZUX | 2/24/16 | pre | BA | 84 | T6.1 | 1.66 | 2 | Q03067 | SGF11_YEAST | 84 |
| 5KGF | 7/27/16 | pre | K | 17 | T1.2 | 0.47 | C | Q12888 | TP53B_HUMAN | 1627 |
| 5KGF | 7/27/16 | pre | N | 17 | T1.1 | 0.22 | C | Q12888 | TP53B_HUMAN | 1627 |
| 5MLU | 5/10/17 | pre | I | 6 | T1.2 | 1.15 | C | P14349 | GAG_FOAMV | 540 |
| 5VEY | 5/17/17 | pre | C | 51 | T1.1 | 2.54 | C | Q8NCN4 | RN169_HUMAN | 27 |
| 5GTC | 6/28/17 | pre | K | 5 | T1.1 | 0.99 | C | Q9QR71 | LANA1_HHV8P | 9 |
| 6NN6 | 2/13/19 | pre | K | 282 | T7.2 | 2.78 | 2 | Q8TEK3 | DOT1L_HUMAN | 282 |
| 6NOG | 2/20/19 | pre | J | 282 | T6.1 | 2.03 | 2 | Q8TEK3 | DOT1L_HUMAN | 282 |
| 6NJ9 | 2/20/19 | pre | K | 278 | T2.1 | 4.87 | 1 | Q8TEK3 | DOT1L_HUMAN | 278 |
| 6NJ9 | 2/20/19 | pre | K | 282 | T6.1 | 2.27 | 2 | Q8TEK3 | DOT1L_HUMAN | 282 |
| 6NJ9 | 2/20/19 | pre | M | 278 | T2.2 | 4.92 | 1 | Q8TEK3 | DOT1L_HUMAN | 278 |
| 6NJ9 | 2/20/19 | pre | M | 282 | T6.2 | 2.24 | 2 | Q8TEK3 | DOT1L_HUMAN | 282 |
| 6NQA | 2/20/19 | pre | F | 278 | T2.2 | 4.27 | 1 | Q8TEK3 | DOT1L_HUMAN | 278 |
| 6NQA | 2/20/19 | pre | F | 282 | T6.2 | 2.11 | 2 | Q8TEK3 | DOT1L_HUMAN | 282 |
| 6J99 | 2/27/19 | pre | L | 282 | T7.2 | 2.32 | 2 | Q8TEK3 | DOT1L_HUMAN | 282 |
| 6N1Z | 3/20/19 | pre | A | 898 | T1.1 | 0.97 | C | Q96P70 | IPO9_HUMAN | 898 |
| 6N1Z | 3/20/19 | pre | A | 896 | T4.1 | 5.99 | 0 | Q96P70 | IPO9_HUMAN | 896 |
| 6O96 | 4/24/19 | pre | K | 281 | T6.2 | 2.2 | 2 | Q8TEK3 | DOT1L_HUMAN | 282 |
| 6JM9 | 5/15/19 | pre | K | 278 | T7.2 | 4.02 | 2 | Q8TEK3 | DOT1L_HUMAN | 282 |
| 6E0P | 5/22/19 | pre | K | 122 | T1.2 | 1.05 | C |  |  |  |
| 6E0P | 5/22/19 | pre | K | 124 | T4.2 | 4.48 | 0 |  |  |  |
| 6E0P | 5/22/19 | pre | L | 122 | T1.1 | 1.1 | C |  |  |  |
| 6E0P | 5/22/19 | pre | L | 124 | T3.1 | 4.53 | 2 |  |  |  |
| 6DZT | 5/22/19 | pre | K | 122 | T1.1 | 0.8 | C |  |  |  |
| 6DZT | 5/22/19 | pre | K | 124 | T4.1 | 4.36 | 0 |  |  |  |
| 6DZT | 5/22/19 | pre | K | 186 | T7.1 | 5.67 | 2 |  |  |  |
| 6DZT | 5/22/19 | pre | L | 122 | T1.2 | 0.7 | C |  |  |  |
| 6DZT | 5/22/19 | pre | L | 124 | T4.2 | 4.5 | 0 |  |  |  |
| 6E0C | 5/22/19 | pre | K | 122 | T1.2 | 0.97 | C |  |  |  |
| 6E0C | 5/22/19 | pre | K | 124 | T4.2 | 4.5 | 0 |  |  |  |
| 6E0C | 5/22/19 | pre | L | 122 | T1.1 | 0.92 | C |  |  |  |
| 6E0C | 5/22/19 | pre | L | 124 | T4.1 | 4.51 | 0 |  |  |  |
| 6OM3 | 7/10/19 | pre | K | 31 | T5.2 | 3.3 | C | P54784 | ORC1_YEAST | 31 |
| 6MUP | 7/24/19 | pre | L | 5 | T1.2 | 1.32 | C | Q03188 | CENPC_HUMAN | 522 |
| 6MUO | 7/24/19 | pre | K | 5 | T1.1 | 1.67 | C | Q03188 | CENPC_HUMAN | 522 |
| 6MUP | 7/24/19 | pre | K | 5 | T1.1 | 0.51 | C | Q03188 | CENPC_HUMAN | 522 |
| 6SEE | 8/14/19 | pre | K | 109 | T5.2 | 2.5 | C | Q03188 | CENPC_HUMAN | 522 |
| 6SE6 | 8/14/19 | pre | K | 109 | T1.1 | 1.01 | C | Q03188 | CENPC_HUMAN | 522 |
| 6SEF | 8/14/19 | pre | K | 109 | T1.1 | 3.53 | C | Q03188 | CENPC_HUMAN | 522 |
| 6QLD | 10/2/19 | pre | A | 6 | T1.2 | 1.09 | C | P35201 | CENPC_YEAST | 289 |
| 6UH5 | 11/20/19 | pre | L | 99 | T2.1 | 2.87 | 1 | Q6CIT4 | SET1_KLULA | 824 |
| 6UH5 | 11/20/19 | pre | L | 103 | T6.1 | 0.99 | 2 | Q6CIT4 | SET1_KLULA | 828 |
| 6VEN | 1/15/20 | pre | N | 179 | T1.1 | 1.27 | C | P38827 | SET1_YEAST | 901 |
| 6VEN | 1/15/20 | pre | N | 182 | T2.1 | 3.01 | 1 | P38827 | SET1_YEAST | 904 |
| 6VEN | 1/15/20 | pre | N | 186 | T6.1 | 2.06 | 2 | P38827 | SET1_YEAST | 908 |
| 6T9L | 1/29/20 | pre | M | 91 | T2.2 | 3.85 | 1 | Q03067 | SGF11_YEAST | 91 |
| 6T9L | 1/29/20 | pre | M | 84 | T6.2 | 2.04 | 2 | Q03067 | SGF11_YEAST | 84 |
| 6LTJ | 2/12/20 | pre | M | 328 | T6.1 | 0.55 | 2 | Q12824 | SNF5_HUMAN | 377 |
| 6LTJ | 2/12/20 | pre | M | 325 | T4.1 | 5.88 | 0 | Q12824 | SNF5_HUMAN | 374 |
| 6LTJ | 2/12/20 | pre | M | 324 | T2.1 | 2.82 | 1 | Q12824 | SNF5_HUMAN | 373 |
| 6LTJ | 2/12/20 | pre | M | 317 | T1.1 | 4.91 | C | Q12824 | SNF5_HUMAN | 366 |
| 6LTJ | 2/12/20 | pre | M | 321 | T5.1 | 2.23 | C | Q12824 | SNF5_HUMAN | 370 |
| 6TDA | 3/18/20 | pre | K | 404 | T7.1 | 0.43 | 2 | Q06168 | SFH1_YEAST | 404 |
| 6TDA | 3/18/20 | pre | K | 401 | T4.1 | 5.5 | 0 | Q06168 | SFH1_YEAST | 401 |
| 6TDA | 3/18/20 | pre | K | 400 | T2.1 | 3.11 | 1 | Q06168 | SFH1_YEAST | 400 |
| 6TDA | 3/18/20 | pre | K | 397 | T1.1 | 1.1 | C | Q06168 | SFH1_YEAST | 397 |
| 6PA7 | 6/17/20 | pre | N | 740 | T7.1 | 1.83 | 2 | Q9UBC3 | DNM3B_HUMAN | 823 |
| 6PA7 | 6/17/20 | pre | N | 743 | T1.1 | 1.65 | C | Q9UBC3 | DNM3B_HUMAN | 826 |
| 7J09 | 9/16/20 | pre | K | 81 | T2.2 | 3.19 | 1 | Q8C6L5 | CGAS_MOUSE | 222 |
| 7J09 | 9/16/20 | pre | K | 100 | T1.2 | 1.51 | C | Q8C6L5 | CGAS_MOUSE | 241 |
| 6X5A | 9/16/20 | pre | K | 87 | T2.1 | 3.37 | 1 | Q8C6L5 | CGAS_MOUSE | 222 |
| 6X5A | 9/16/20 | pre | K | 106 | T1.1 | 1.04 | C | Q8C6L5 | CGAS_MOUSE | 241 |
| 7J0A | 9/16/20 | pre | K | 81 | T2.2 | 3.67 | 1 | Q8C6L5 | CGAS_MOUSE | 222 |
| 7J0A | 9/16/20 | pre | K | 100 | T1.2 | 1.83 | C | Q8C6L5 | CGAS_MOUSE | 241 |
| 6XJD | 9/16/20 | pre | K | 87 | T2.1 | 3.65 | 1 | Q8C6L5 | CGAS_MOUSE | 222 |
| 6XJD | 9/16/20 | pre | K | 106 | T1.1 | 0.86 | C | Q8C6L5 | CGAS_MOUSE | 241 |
| 6XJD | 9/16/20 | pre | L | 87 | T2.2 | 3.32 | 1 | Q8C6L5 | CGAS_MOUSE | 222 |
| 6XJD | 9/16/20 | pre | L | 106 | T1.2 | 0.8 | C | Q8C6L5 | CGAS_MOUSE | 241 |
| 6X59 | 9/16/20 | pre | K | 87 | T2.1 | 2.67 | 1 | Q8C6L5 | CGAS_MOUSE | 222 |
| 6X59 | 9/16/20 | pre | K | 106 | T1.1 | 1.07 | C | Q8C6L5 | CGAS_MOUSE | 241 |
| 7C0M | 9/16/20 | pre | K | 88 | T2.4 | 2.69 | 1 | Q8N884 | CGAS_HUMAN | 236 |
| 7C0M | 9/16/20 | pre | K | 107 | T1.4 | 1.02 | C | Q8N884 | CGAS_HUMAN | 255 |
| 7C0M | 9/16/20 | pre | V | 88 | T2.2 | 2.69 | 1 | Q8N884 | CGAS_HUMAN | 236 |
| 7C0M | 9/16/20 | pre | V | 107 | T1.2 | 1.02 | C | Q8N884 | CGAS_HUMAN | 255 |
| 6Y5D | 9/23/20 | pre | K | 76 | T2.3 | 3.77 | 1 | Q8N884 | CGAS_HUMAN | 236 |
| 6Y5D | 9/23/20 | pre | K | 95 | T1.3 | 1.65 | C | Q8N884 | CGAS_HUMAN | 255 |
| 6Y5D | 9/23/20 | pre | L | 76 | T2.1 | 3.94 | 1 | Q8N884 | CGAS_HUMAN | 236 |
| 6Y5D | 9/23/20 | pre | L | 95 | T1.1 | 1.37 | C | Q8N884 | CGAS_HUMAN | 255 |
| 6Y5E | 9/23/20 | pre | K | 76 | T2.1 | 2.74 | 1 | Q8N884 | CGAS_HUMAN | 236 |
| 6Y5E | 9/23/20 | pre | K | 95 | T1.1 | 1.07 | C | Q8N884 | CGAS_HUMAN | 255 |
| 7A08 | 9/23/20 | pre | A | 85 | T2.2 | 2.86 | 1 | Q8C6L5 | CGAS_MOUSE | 222 |
| 7A08 | 9/23/20 | pre | A | 104 | T1.2 | 0.76 | C | Q8C6L5 | CGAS_MOUSE | 241 |
| 7CCR | 10/7/20 | pre | K | 80 | T2.4 | 2.77 | 1 | Q8N884 | CGAS_HUMAN | 236 |
| 7CCR | 10/7/20 | pre | K | 99 | T1.4 | 1.71 | C | Q8N884 | CGAS_HUMAN | 255 |
| 7CCR | 10/7/20 | pre | V | 80 | T2.2 | 2.77 | 1 | Q8N884 | CGAS_HUMAN | 236 |
| 7CCR | 10/7/20 | pre | V | 99 | T1.2 | 1.71 | C | Q8N884 | CGAS_HUMAN | 255 |
| 7CCQ | 10/7/20 | pre | I | 80 | T2.2 | 2.47 | 1 | Q8N884 | CGAS_HUMAN | 236 |
| 7CCQ | 10/7/20 | pre | I | 99 | T1.2 | 1.75 | C | Q8N884 | CGAS_HUMAN | 255 |
| 7K63 | 11/25/20 | pre | K | 252 | T1.2 | 0.73 | C |  |  |  |
| 7K63 | 11/25/20 | pre | K | 254 | T4.2 | 4.25 | 0 |  |  |  |
| 7K63 | 11/25/20 | pre | L | 252 | T1.1 | 0.62 | C |  |  |  |
| 7K63 | 11/25/20 | pre | L | 254 | T4.1 | 4.21 | 0 |  |  |  |
| 7K60 | 11/25/20 | pre | K | 252 | T1.2 | 1.17 | C |  |  |  |
| 7K60 | 11/25/20 | pre | K | 254 | T4.2 | 4.49 | 0 |  |  |  |
| 7K60 | 11/25/20 | pre | L | 252 | T1.1 | 1.06 | C |  |  |  |
| 7K60 | 11/25/20 | pre | L | 254 | T4.1 | 4.58 | 0 |  |  |  |
| 7K61 | 11/25/20 | pre | K | 252 | T1.2 | 0.84 | C |  |  |  |
| 7K61 | 11/25/20 | pre | K | 254 | T4.2 | 4.48 | 0 |  |  |  |
| 7K61 | 11/25/20 | pre | L | 252 | T1.1 | 0.72 | C |  |  |  |
| 7K61 | 11/25/20 | pre | L | 254 | T4.1 | 4.58 | 0 |  |  |  |
| 7K5Y | 11/25/20 | pre | K | 252 | T1.2 | 1.01 | C |  |  |  |
| 7K5Y | 11/25/20 | pre | K | 254 | T4.2 | 4.49 | 0 |  |  |  |
| 7K5Y | 11/25/20 | pre | L | 252 | T1.1 | 1.1 | C |  |  |  |
| 7K5Y | 11/25/20 | pre | L | 254 | T4.1 | 4.54 | 0 |  |  |  |
| 7K5X | 11/25/20 | pre | K | 252 | T1.2 | 0.9 | C |  |  |  |
| 7K5X | 11/25/20 | pre | K | 254 | T4.2 | 4.46 | 0 |  |  |  |
| 7K5X | 11/25/20 | pre | L | 252 | T1.1 | 0.87 | C |  |  |  |
| 7K5X | 11/25/20 | pre | L | 254 | T4.1 | 4.75 | 0 |  |  |  |
| 7D20 | 2/10/21 | pre | J | 192 | T1.1 | 0.74 | C | Q9NQR1 | KMT5A_HUMAN | 233 |
| 7D1Z | 2/10/21 | pre | J | 192 | T1.1 | 0.83 | C | Q9NQR1 | KMT5A_HUMAN | 233 |
| 7BXT | 2/10/21 | pre | K | 17 | T1.2 | 2.38 | C | O57392 | O57392_CHICK | 659 |
| 7BXT | 2/10/21 | pre | L | 17 | T1.1 | 1.47 | C | O57392 | O57392_CHICK | 659 |
| 7BXT | 2/10/21 | pre | K | 19 | T2.2 | 4.14 | 1 | O57392 | O57392_CHICK | 661 |
| 7BXT | 2/10/21 | pre | L | 19 | T2.1 | 3.48 | 1 | O57392 | O57392_CHICK | 661 |
| 7BY0 | 2/10/21 | pre | K | 455 | T6.2 | 2.06 | 2 | O57393 | O57393_CHICK | 644 |
| 7BY0 | 2/10/21 | pre | K | 459 | T5.2 | 1.96 | C | O57393 | O57393_CHICK | 648 |
| 7BY0 | 2/10/21 | pre | K | 461 | T2.2 | 4.92 | 1 | O57393 | O57393_CHICK | 650 |
| 7BY0 | 2/10/21 | pre | L | 455 | T6.1 | 2.12 | 2 | O57393 | O57393_CHICK | 644 |
| 7BY0 | 2/10/21 | pre | L | 459 | T5.1 | 2.61 | C | O57393 | O57393_CHICK | 648 |
| 7BY0 | 2/10/21 | pre | L | 461 | T2.1 | 5.6 | 1 | O57393 | O57393_CHICK | 650 |
| 7JZV | 2/17/21 | pre | A | 72 | T1.2 | 1.5 | C | P38398 | BRCA1_HUMAN | 71 |
| 7K78 | 3/31/21 | pre | K | 122 | T1.2 | 0.69 | C |  |  |  |
| 7K78 | 3/31/21 | pre | K | 124 | T4.2 | 4.53 | 0 |  |  |  |
| 7K78 | 3/31/21 | pre | L | 122 | T1.1 | 1.17 | C |  |  |  |
| 7K78 | 3/31/21 | pre | L | 124 | T4.1 | 4.75 | 0 |  |  |  |
| 7BWD | 4/14/21 | pre | A | 282 | T7.2 | 2.38 | 2 | Q8TEK3 | DOT1L_HUMAN | 282 |
| 7LYC | 6/16/21 | pre | J | 301 | T1.2 | 0.52 | C | Q99728 | BARD1_HUMAN | 705 |
| 7E8I | 6/30/21 | pre | K | 284 | T1.2 | 1.39 | C | Q99728 | BARD1_HUMAN | 705 |
| 7ON1 | 7/7/21 | pre | A | 289 | T1.2 | 1.3 | C | P35201 | CENPC_YEAST | 289 |
| 7ON1 | 7/7/21 | pre | L | 289 | T1.1 | 1.77 | C | P35201 | CENPC_YEAST | 289 |
| 7ENN | 7/14/21 | pre | A | 614 | T1.2 | 0.96 | C | Q86WJ1 | CHD1L_HUMAN | 614 |
| 7LYB | 7/28/21 | pre | I | 95 | T1.1 | 1.22 | C | P38398 | BRCA1_HUMAN | 71 |
| 7EGP | 1/12/22 | post | B | 676 | T3.1 | 0.94 | 2 | P18480 | SNF5_YEAST | 676 |
| 7EG6 | 1/12/22 | post | A | 676 | T6.1 | 1.37 | 2 | P18480 | SNF5_YEAST | 676 |
| 7EGP | 1/12/22 | post | B | 673 | T4.1 | 4.09 | 0 | P18480 | SNF5_YEAST | 673 |
| 7EG6 | 1/12/22 | post | A | 673 | T4.1 | 4.89 | 0 | P18480 | SNF5_YEAST | 673 |
| 7EG6 | 1/12/22 | post | A | 672 | T2.1 | 3.36 | 1 | P18480 | SNF5_YEAST | 672 |
| 7EGP | 1/12/22 | post | B | 672 | T2.1 | 3.57 | 1 | P18480 | SNF5_YEAST | 672 |
| 7EG6 | 1/12/22 | post | A | 669 | T1.1 | 0.9 | C | P18480 | SNF5_YEAST | 669 |
| 7EGP | 1/12/22 | post | B | 669 | T1.1 | 1.79 | C | P18480 | SNF5_YEAST | 669 |
| 7TN2 | 3/2/22 | post | K | 11 | T6.1 | 2.35 | 2 | P32657 | CHD1_YEAST | 126 |
| 7TN2 | 3/2/22 | post | K | 15 | T1.1 | 1.04 | C | P32657 | CHD1_YEAST | 130 |
| 7XCT | 4/20/22 | post | E | 274 | T2.2 | 4.88 | 1 | Q8TEK3 | DOT1L_HUMAN | 278 |
| 7XCT | 4/20/22 | post | E | 278 | T6.2 | 1.62 | 2 | Q8TEK3 | DOT1L_HUMAN | 282 |
| 7XCT | 4/20/22 | post | M | 274 | T2.1 | 4.88 | 1 | Q8TEK3 | DOT1L_HUMAN | 278 |
| 7XCT | 4/20/22 | post | M | 278 | T6.1 | 1.8 | 2 | Q8TEK3 | DOT1L_HUMAN | 282 |
| 7XCR | 4/20/22 | post | E | 274 | T2.2 | 5.46 | 1 | Q8TEK3 | DOT1L_HUMAN | 278 |
| 7XCR | 4/20/22 | post | E | 278 | T6.2 | 1.35 | 2 | Q8TEK3 | DOT1L_HUMAN | 282 |
| 7R5R | 4/27/22 | post | K | 521 | T1.1 | 1.51 | C | Q03188 | CENPC_HUMAN | 522 |
| 7R5R | 4/27/22 | post | L | 521 | T1.2 | 1.4 | C | Q03188 | CENPC_HUMAN | 522 |
| 7VDV | 5/18/22 | post | Q | 377 | T6.1 | 1.63 | 2 | Q12824 | SNF5_HUMAN | 377 |
| 7VDV | 5/18/22 | post | J | 1183 | T2.2 | 4.1 | 1 | P51532 | SMCA4_HUMAN | 1374 |
| 7VDV | 5/18/22 | post | Q | 373 | T2.1 | 3.47 | 1 | Q12824 | SNF5_HUMAN | 373 |
| 7VDT | 5/18/22 | post | A | 1183 | T2.2 | 3.1 | 1 | P51532 | SMCA4_HUMAN | 1374 |
| 7VDV | 5/18/22 | post | J | 1178 | T6.2 | 2.73 | 2 | P51532 | SMCA4_HUMAN | 1369 |
| 7VDT | 5/18/22 | post | A | 1178 | T6.2 | 1.37 | 2 | P51532 | SMCA4_HUMAN | 1369 |
| 7YYH | 5/18/22 | post | U | 522 | T1.1 | 1.55 | C | Q03188 | CENPC_HUMAN | 522 |
| 7YWX | 5/18/22 | post | Z | 522 | T1.1 | 1.51 | C | Q03188 | CENPC_HUMAN | 522 |
| 7YYH | 5/18/22 | post | V | 522 | T1.2 | 1.28 | C | Q03188 | CENPC_HUMAN | 522 |
| 7VDV | 5/18/22 | post | J | 1181 | T1.2 | 2.04 | C | P51532 | SMCA4_HUMAN | 1372 |
| 7YWX | 5/18/22 | post | AA | 522 | T1.2 | 1.4 | C | Q03188 | CENPC_HUMAN | 522 |
| 7VDT | 5/18/22 | post | A | 1181 | T1.2 | 2.05 | C | P51532 | SMCA4_HUMAN | 1372 |
| 7VDV | 5/18/22 | post | Q | 370 | T5.1 | 2.35 | C | Q12824 | SNF5_HUMAN | 370 |
| 7PII | 5/25/22 | post | K | 522 | T1.1 | 1.55 | C | Q03188 | CENPC_HUMAN | 522 |
| 7PII | 5/25/22 | post | L | 522 | T1.2 | 1.28 | C | Q03188 | CENPC_HUMAN | 522 |
| 7XPX | 6/1/22 | post | K | 61 | T1.2 | 1.06 | C | Q9NQR1 | KMT5A_HUMAN | 233 |
| 7Y7I | 7/13/22 | post | K | 8 | T7.2 | 4.67 | 2 |  |  |  |
| 7Y7I | 7/13/22 | post | K | 11 | T1.2 | 1.09 | C |  |  |  |
| 7Y7I | 7/13/22 | post | L | 11 | T1.1 | 1.3 | C |  |  |  |
| 7VVZ | 8/10/22 | post | O | 4 | T1.1 | 0.79 | C | P43572 | EPL1_YEAST | 58 |
| 7VVU | 8/10/22 | post | O | 58 | T1.1 | 0.79 | C | P43572 | EPL1_YEAST | 58 |
| 7YQK | 8/17/22 | post | L | 15 | T5.1 | 1.55 | C | Q12888 | TP53B_HUMAN | 1627 |
| 7XSZ | 9/7/22 | post | R | 410 | T8.1 | 5.38 | 2 | C4R370 | C4R370_KOMPG | 407 |
| 7E9C | 9/7/22 | post | K | 28 | T7.2 | 3 | 2 | P54784 | ORC1_YEAST | 29 |
| 7E9C | 9/7/22 | post | K | 30 | T1.2 | 1.96 | C | P54784 | ORC1_YEAST | 31 |
| 7E9F | 9/7/22 | post | K | 29 | T6.2 | 5.25 | 2 | P54784 | ORC1_YEAST | 29 |
| 7E9F | 9/7/22 | post | K | 31 | T1.2 | 1.81 | C | P54784 | ORC1_YEAST | 31 |
| 7E9F | 9/7/22 | post | K | 33 | T1.2 | 4.95 | C | P54784 | ORC1_YEAST | 33 |
| 7E9F | 9/7/22 | post | L | 29 | T6.1 | 5.88 | 2 | P54784 | ORC1_YEAST | 29 |
| 7E9F | 9/7/22 | post | L | 31 | T1.1 | 1.91 | C | P54784 | ORC1_YEAST | 31 |
| 7E9F | 9/7/22 | post | L | 33 | T1.1 | 5.26 | C | P54784 | ORC1_YEAST | 33 |
| 7Y8R | 12/7/22 | post | N | 377 | T6.1 | 1.74 | 2 | Q12824 | SNF5_HUMAN | 377 |
| 7Y8R | 12/7/22 | post | N | 374 | T4.1 | 4.07 | 0 | Q12824 | SNF5_HUMAN | 374 |
| 7Y8R | 12/7/22 | post | N | 373 | T2.1 | 3.81 | 1 | Q12824 | SNF5_HUMAN | 373 |
| 7Y8R | 12/7/22 | post | N | 370 | T5.1 | 1.04 | C | Q12824 | SNF5_HUMAN | 370 |
| 8AV6 | 12/14/22 | post | J | 501 | T5.1 | 2.11 | C | G0S589 | G0S589_CHATD | 598 |
| 8H1T | 2/1/23 | post | L | 59 | T5.2 | 2.64 | C | Q92560 | BAP1_HUMAN | 59 |
| 8F86 | 4/5/23 | post | K | 172 | T6.2 | 3.07 | 2 | Q8N6T7 | SIR6_HUMAN | 172 |
| 8F86 | 4/5/23 | post | K | 175 | T1.2 | 1.18 | C | Q8N6T7 | SIR6_HUMAN | 175 |
| 8F86 | 4/5/23 | post | K | 178 | T5.2 | 5.89 | C | Q8N6T7 | SIR6_HUMAN | 178 |
| 8GRM | 4/12/23 | post | J | 83 | T1.1 | 1.45 | C | Q99496 | RING2_HUMAN | 98 |
| 8GRQ | 4/12/23 | post | K | 66 | T4.2 | 2.54 | 0 | P38398 | BRCA1_HUMAN | 71 |
| 8G57 | 4/26/23 | post | A | 174 | T6.1 | 2.06 | 2 | Q8N6T7 | SIR6_HUMAN | 172 |
| 8G57 | 4/26/23 | post | A | 177 | T1.1 | 1.06 | C | Q8N6T7 | SIR6_HUMAN | 175 |
| 8G57 | 4/26/23 | post | A | 180 | T5.1 | 3.94 | C | Q8N6T7 | SIR6_HUMAN | 178 |
| 8DK5 | 6/28/23 | post | I | 122 | T1.1 | 0.88 | C |  |  |  |
| 8DK5 | 6/28/23 | post | I | 124 | T4.1 | 4.68 | 0 |  |  |  |
| 8DK5 | 6/28/23 | post | J | 122 | T1.2 | 1.04 | C |  |  |  |
| 8DK5 | 6/28/23 | post | J | 124 | T4.2 | 4.49 | 0 |  |  |  |
| 8SPU | 6/28/23 | post | L | 122 | T1.1 | 0.88 | C |  |  |  |
| 8SPU | 6/28/23 | post | L | 124 | T4.1 | 4.68 | 0 |  |  |  |
| 8SPU | 6/28/23 | post | M | 122 | T1.2 | 1.04 | C |  |  |  |
| 8SPU | 6/28/23 | post | M | 124 | T4.2 | 4.49 | 0 |  |  |  |
| 7U0J | 6/28/23 | post | K | 122 | T1.1 | 0.88 | C |  |  |  |
| 7U0J | 6/28/23 | post | K | 124 | T4.1 | 4.68 | 0 |  |  |  |
| 7U0J | 6/28/23 | post | L | 122 | T1.2 | 1.04 | C |  |  |  |
| 7U0J | 6/28/23 | post | L | 124 | T4.2 | 4.49 | 0 |  |  |  |
| 7U0I | 6/28/23 | post | M | 122 | T1.1 | 0.85 | C |  |  |  |
| 7U0I | 6/28/23 | post | M | 124 | T4.1 | 4.64 | 0 |  |  |  |
| 7U0I | 6/28/23 | post | N | 122 | T1.2 | 1.01 | C |  |  |  |
| 7U0I | 6/28/23 | post | N | 124 | T4.2 | 4.61 | 0 |  |  |  |
| 7U0G | 6/28/23 | post | N | 122 | T1.1 | 0.94 | C |  |  |  |
| 7U0G | 6/28/23 | post | N | 124 | T4.1 | 4.7 | 0 |  |  |  |
| 7U0G | 6/28/23 | post | O | 122 | T1.2 | 0.8 | C |  |  |  |
| 7U0G | 6/28/23 | post | O | 124 | T4.2 | 4.51 | 0 |  |  |  |
| 7U0G | 6/28/23 | post | O | 186 | T7.2 | 5.94 | 2 |  |  |  |
| 8SPS | 6/28/23 | post | L | 122 | T1.1 | 0.88 | C |  |  |  |
| 8SPS | 6/28/23 | post | L | 124 | T4.1 | 4.68 | 0 |  |  |  |
| 8SPS | 6/28/23 | post | M | 122 | T1.2 | 1.04 | C |  |  |  |
| 8SPS | 6/28/23 | post | M | 124 | T4.2 | 4.49 | 0 |  |  |  |
| 8FOX | 7/26/23 | post | A | 864 | T1.1 | 0.75 | C | P53067 | IMB5_YEAST | 864 |
| 8F1E | 7/26/23 | post | A | 864 | T1.1 | 2.56 | C | P53067 | IMB5_YEAST | 864 |
| 8OF4 | 8/9/23 | post | K | 172 | T6.2 | 2.62 | 2 | Q8N6T7 | SIR6_HUMAN | 172 |
| 8OF4 | 8/9/23 | post | K | 175 | T1.2 | 1.6 | C | Q8N6T7 | SIR6_HUMAN | 175 |
| 8OF4 | 8/9/23 | post | K | 178 | T5.2 | 4.24 | C | Q8N6T7 | SIR6_HUMAN | 178 |
| 7YRD | 8/16/23 | post | K | 273 | T2.2 | 2.71 | 1 | Q4FZB7 | KMT5B_HUMAN | 357 |
| 7YRD | 8/16/23 | post | K | 268 | T1.2 | 1.26 | C | Q4FZB7 | KMT5B_HUMAN | 352 |
| 8SVF | 8/30/23 | post | K | 56 | T4.2 | 3.26 | 0 | Q92560 | BAP1_HUMAN | 56 |
| 8T9F | 9/6/23 | post | A | 352 | T1.1 | 0.88 | C | Q4FZB7 | KMT5B_HUMAN | 352 |
| 8IEG | 9/6/23 | post | K | 43 | T1.2 | 0.68 | C | Q07457 | BRE1_YEAST | 679 |
| 8IEG | 9/6/23 | post | K | 45 | T2.2 | 2.08 | 1 | Q07457 | BRE1_YEAST | 681 |
| 8IEJ | 9/6/23 | post | L | 39 | T1.2 | 0.84 | C | Q5VTR2 | BRE1A_HUMAN | 953 |
| 8IEJ | 9/6/23 | post | L | 41 | T2.2 | 2.91 | 1 | Q5VTR2 | BRE1A_HUMAN | 955 |
| 8T9F | 9/6/23 | post | A | 357 | T2.1 | 3.16 | 1 | Q4FZB7 | KMT5B_HUMAN | 357 |
| 8T9H | 9/13/23 | post | K | 358 | T2.2 | 3.67 | 1 | Q4FZB7 | KMT5B_HUMAN | 357 |
| 8HR1 | 9/13/23 | post | I | 8 | T1.2 | 1.39 | C | Q16384 | SSX1_HUMAN | 169 |
| 8HR1 | 9/13/23 | post | I | 6 | T6.2 | 3.15 | 2 | Q16384 | SSX1_HUMAN | 167 |
| 8T9H | 9/13/23 | post | K | 353 | T5.2 | 1.57 | C | Q4FZB7 | KMT5B_HUMAN | 352 |
| 7X3T | 9/20/23 | post | K | 614 | T4.4 | 5.88 | 0 | P43596 | IOC3_YEAST | 739 |
| 7X3V | 9/20/23 | post | K | 613 | T2.2 | 2.84 | 1 | P43596 | IOC3_YEAST | 738 |
| 7X3V | 9/20/23 | post | K | 610 | T1.2 | 1.22 | C | P43596 | IOC3_YEAST | 735 |
| 7X3T | 9/20/23 | post | K | 610 | T5.4 | 2.83 | C | P43596 | IOC3_YEAST | 735 |
| 8GUJ | 9/20/23 | post | L | 979 | T1.2 | 0.63 | C | O75150 | BRE1B_HUMAN | 979 |
| 8GUJ | 9/20/23 | post | L | 981 | T2.2 | 2.28 | 1 | O75150 | BRE1B_HUMAN | 981 |
| 8GUI | 9/20/23 | post | K | 953 | T1.2 | 0.78 | C | Q5VTR2 | BRE1A_HUMAN | 953 |
| 8GUI | 9/20/23 | post | K | 955 | T2.2 | 2.57 | 1 | Q5VTR2 | BRE1A_HUMAN | 955 |
| 8HQY | 9/27/23 | post | I | 6 | T6.2 | 5.43 | 2 | Q16385 | SSX2_HUMAN | 167 |
| 8HQY | 9/27/23 | post | I | 8 | T1.2 | 0.92 | C | Q16385 | SSX2_HUMAN | 169 |
| 8T3W | 10/18/23 | post | K | 89 | T1.1 | 1.73 | C | Q07457 | BRE1_YEAST | 679 |
| 8T3W | 10/18/23 | post | K | 91 | T2.1 | 3.55 | 1 | Q07457 | BRE1_YEAST | 681 |
| 8T3Y | 10/18/23 | post | K | 89 | T1.1 | 1.94 | C | Q07457 | BRE1_YEAST | 679 |
| 8T3Y | 10/18/23 | post | K | 91 | T2.1 | 3.31 | 1 | Q07457 | BRE1_YEAST | 681 |
| 8T3T | 10/18/23 | post | L | 89 | T1.1 | 1.36 | C | Q07457 | BRE1_YEAST | 679 |
| 8T3T | 10/18/23 | post | L | 91 | T2.1 | 2.84 | 1 | Q07457 | BRE1_YEAST | 681 |
| 8EVJ | 11/1/23 | post | K | 122 | T1.1 | 1.86 | C |  |  |  |
| 8EVJ | 11/1/23 | post | K | 124 | T4.1 | 5.74 | 0 |  |  |  |
| 8EVJ | 11/1/23 | post | L | 122 | T1.2 | 0.95 | C |  |  |  |
| 8EVJ | 11/1/23 | post | L | 124 | T3.2 | 4.71 | 2 |  |  |  |
| 8EVH | 11/1/23 | post | K | 122 | T1.1 | 0.86 | C |  |  |  |
| 8EVH | 11/1/23 | post | K | 124 | T4.1 | 4.78 | 0 |  |  |  |
| 8EVH | 11/1/23 | post | L | 122 | T1.2 | 0.92 | C |  |  |  |
| 8EVH | 11/1/23 | post | L | 124 | T4.2 | 4.74 | 0 |  |  |  |
| 8EVG | 11/1/23 | post | K | 122 | T1.1 | 0.9 | C |  |  |  |
| 8EVG | 11/1/23 | post | K | 124 | T4.1 | 4.76 | 0 |  |  |  |
| 8EVG | 11/1/23 | post | L | 122 | T1.2 | 1.04 | C |  |  |  |
| 8EVG | 11/1/23 | post | L | 124 | T4.2 | 4.6 | 0 |  |  |  |
| 8SYP | 11/1/23 | post | K | 122 | T1.1 | 0.88 | C |  |  |  |
| 8SYP | 11/1/23 | post | K | 124 | T4.1 | 4.76 | 0 |  |  |  |
| 8SYP | 11/1/23 | post | L | 122 | T1.2 | 0.84 | C |  |  |  |
| 8SYP | 11/1/23 | post | L | 124 | T4.2 | 4.61 | 0 |  |  |  |
| 8EVI | 11/1/23 | post | K | 122 | T1.1 | 0.86 | C |  |  |  |
| 8EVI | 11/1/23 | post | K | 124 | T4.1 | 4.78 | 0 |  |  |  |
| 8EVI | 11/1/23 | post | L | 122 | T1.2 | 0.92 | C |  |  |  |
| 8EVI | 11/1/23 | post | L | 124 | T4.2 | 4.74 | 0 |  |  |  |
| 8JHF | 11/15/23 | post | A | 357 | T2.1 | 3.96 | 1 | Q4FZB7 | KMT5B_HUMAN | 357 |
| 8JHF | 11/15/23 | post | A | 352 | T1.1 | 1.76 | C | Q4FZB7 | KMT5B_HUMAN | 352 |
| 8JHG | 11/22/23 | post | A | 357 | T2.1 | 3.72 | 1 | Q4FZB7 | KMT5B_HUMAN | 357 |
| 8JHG | 11/22/23 | post | A | 352 | T1.1 | 1.43 | C | Q4FZB7 | KMT5B_HUMAN | 352 |
| 8UX1 | 11/22/23 | post | L | 216 | T1.2 | 1.12 | C | P18754 | RCC1_HUMAN | 217 |
| 8UX1 | 11/22/23 | post | L | 221 | T3.2 | 5.81 | 2 | P18754 | RCC1_HUMAN | 222 |
| 7YRG | 12/13/23 | post | K | 273 | T2.2 | 2.71 | 1 | Q4FZB7 | KMT5B_HUMAN | 357 |
| 7YRG | 12/13/23 | post | L | 273 | T2.1 | 1.67 | 1 | Q4FZB7 | KMT5B_HUMAN | 357 |
| 7YRG | 12/13/23 | post | K | 268 | T1.2 | 1.26 | C | Q4FZB7 | KMT5B_HUMAN | 352 |
| 7YRG | 12/13/23 | post | L | 268 | T1.1 | 0.59 | C | Q4FZB7 | KMT5B_HUMAN | 352 |
| 8SN2 | 1/17/24 | post | K | 67 | T3.2 | 2.44 | 2 | Q8IYW5 | RN168_HUMAN | 57 |
| 8SN2 | 1/17/24 | post | K | 73 | T1.2 | 1.36 | C | Q8IYW5 | RN168_HUMAN | 63 |
| 8SN2 | 1/17/24 | post | K | 77 | T5.2 | 5.76 | C | Q8IYW5 | RN168_HUMAN | 67 |
| 8SN4 | 1/17/24 | post | K | 67 | T6.2 | 2.37 | 2 | Q8IYW5 | RN168_HUMAN | 57 |
| 8SN4 | 1/17/24 | post | K | 73 | T1.2 | 1.9 | C | Q8IYW5 | RN168_HUMAN | 63 |
| 8SN5 | 1/17/24 | post | K | 67 | T6.2 | 2.28 | 2 | Q8IYW5 | RN168_HUMAN | 57 |
| 8SN5 | 1/17/24 | post | K | 73 | T1.2 | 2.01 | C | Q8IYW5 | RN168_HUMAN | 63 |
| 8SMW | 1/17/24 | post | K | 67 | T6.2 | 2.13 | 2 | Q8IYW5 | RN168_HUMAN | 57 |
| 8SMW | 1/17/24 | post | K | 73 | T5.2 | 1.85 | C | Q8IYW5 | RN168_HUMAN | 63 |
| 8SN6 | 1/17/24 | post | K | 67 | T6.2 | 1.72 | 2 | Q8IYW5 | RN168_HUMAN | 57 |
| 8SN6 | 1/17/24 | post | K | 73 | T1.2 | 1.84 | C | Q8IYW5 | RN168_HUMAN | 63 |
| 8SN6 | 1/17/24 | post | K | 77 | T5.2 | 5.39 | C | Q8IYW5 | RN168_HUMAN | 67 |
| 8SN1 | 1/17/24 | post | K | 67 | T6.2 | 1.99 | 2 | Q8IYW5 | RN168_HUMAN | 57 |
| 8SN1 | 1/17/24 | post | K | 73 | T1.2 | 1.9 | C | Q8IYW5 | RN168_HUMAN | 63 |
| 8SN0 | 1/17/24 | post | K | 67 | T6.2 | 2.41 | 2 | Q8IYW5 | RN168_HUMAN | 57 |
| 8SN0 | 1/17/24 | post | K | 73 | T1.2 | 1.34 | C | Q8IYW5 | RN168_HUMAN | 63 |
| 8SNA | 1/17/24 | post | K | 67 | T6.2 | 2.04 | 2 | Q8IYW5 | RN168_HUMAN | 57 |
| 8SNA | 1/17/24 | post | K | 73 | T1.2 | 1.49 | C | Q8IYW5 | RN168_HUMAN | 63 |
| 8SN3 | 1/17/24 | post | K | 67 | T6.2 | 2.01 | 2 | Q8IYW5 | RN168_HUMAN | 57 |
| 8SN3 | 1/17/24 | post | K | 73 | T1.2 | 1.5 | C | Q8IYW5 | RN168_HUMAN | 63 |
| 8UPF | 1/17/24 | post | I | 59 | T3.1 | 3.16 | 2 | Q8IYW5 | RN168_HUMAN | 57 |
| 8UPF | 1/17/24 | post | I | 65 | T1.1 | 1.38 | C | Q8IYW5 | RN168_HUMAN | 63 |
| 8UPF | 1/17/24 | post | I | 69 | T1.1 | 5.91 | C | Q8IYW5 | RN168_HUMAN | 67 |
| 8U14 | 1/17/24 | post | I | 57 | T3.1 | 2.46 | 2 | Q8IYW5 | RN168_HUMAN | 57 |
| 8U14 | 1/17/24 | post | I | 63 | T1.1 | 2.19 | C | Q8IYW5 | RN168_HUMAN | 63 |
| 8SN9 | 1/17/24 | post | K | 67 | T3.2 | 2.52 | 2 | Q8IYW5 | RN168_HUMAN | 57 |
| 8SN9 | 1/17/24 | post | K | 73 | T1.2 | 1.5 | C | Q8IYW5 | RN168_HUMAN | 63 |
| 8SMX | 1/17/24 | post | K | 67 | T3.2 | 2.48 | 2 | Q8IYW5 | RN168_HUMAN | 57 |
| 8SMX | 1/17/24 | post | K | 73 | T1.2 | 1.35 | C | Q8IYW5 | RN168_HUMAN | 63 |
| 8SMZ | 1/17/24 | post | K | 67 | T6.2 | 1.67 | 2 | Q8IYW5 | RN168_HUMAN | 57 |
| 8SMZ | 1/17/24 | post | K | 73 | T5.2 | 2.39 | C | Q8IYW5 | RN168_HUMAN | 63 |
| 8SN7 | 1/17/24 | post | K | 67 | T6.2 | 2.05 | 2 | Q8IYW5 | RN168_HUMAN | 57 |
| 8SN7 | 1/17/24 | post | K | 73 | T1.2 | 1.41 | C | Q8IYW5 | RN168_HUMAN | 63 |
| 8SN8 | 1/17/24 | post | K | 67 | T6.2 | 2.15 | 2 | Q8IYW5 | RN168_HUMAN | 57 |
| 8SN8 | 1/17/24 | post | K | 73 | T1.2 | 1.96 | C | Q8IYW5 | RN168_HUMAN | 63 |
| 8SMY | 1/17/24 | post | K | 67 | T6.2 | 1.88 | 2 | Q8IYW5 | RN168_HUMAN | 57 |
| 8SMY | 1/17/24 | post | K | 73 | T1.2 | 1.34 | C | Q8IYW5 | RN168_HUMAN | 63 |
| 8U13 | 1/17/24 | post | I | 57 | T7.1 | 5.12 | 2 | Q8IYW5 | RN168_HUMAN | 57 |
| 8U13 | 1/17/24 | post | I | 63 | T1.1 | 1.77 | C | Q8IYW5 | RN168_HUMAN | 63 |
| 8OL1 | 2/14/24 | post | K | 76 | T2.1 | 2.76 | 1 | Q8N884 | CGAS_HUMAN | 236 |
| 8OL1 | 2/14/24 | post | K | 95 | T5.1 | 0.59 | C | Q8N884 | CGAS_HUMAN | 255 |
| 8X1C | 3/6/24 | post | J | 134 | T1.1 | 2.05 | C | Q15906 | VPS72_HUMAN | 134 |
| 8X19 | 3/6/24 | post | J | 134 | T1.1 | 1.26 | C | Q15906 | VPS72_HUMAN | 134 |
| 8X19 | 3/6/24 | post | L | 9 | T8.2 | 3.11 | 2 | O43257 | ZNHI1_HUMAN | 9 |
| 8X19 | 3/6/24 | post | L | 17 | T1.2 | 5.2 | C | O43257 | ZNHI1_HUMAN | 17 |
| 8IHT | 3/20/24 | post | K | 61 | T5.2 | 2.86 | C | Q04779 | RCO1_YEAST | 61 |
| 8IHM | 3/20/24 | post | K | 12 | T5.2 | 2.86 | C | Q04779 | RCO1_YEAST | 61 |
| 8Q36 | 3/27/24 | post | A | 6 | T1.1 | 0.96 | C |  |  |  |
| 8Q3E | 3/27/24 | post | A | 6 | T1.1 | 0.86 | C |  |  |  |
| 8PP7 | 3/27/24 | post | B | 102 | T1.2 | 1.32 | C | Q99496 | RING2_HUMAN | 98 |
| 8PP7 | 3/27/24 | post | D | 102 | T1.1 | 1.43 | C | Q99496 | RING2_HUMAN | 98 |

**Table 2.** Final acidic patch interactor hit list as determined by in silico screening with AlphaFold-Multimer and AlphaFold 3.

| UniProt ID | UniProt Entry Name | Gene | Anchor count | Anchor indices | In PDB | Detected by MS |
| --- | --- | --- | --- | --- | --- | --- |
| Q03188 | CENPC_HUMAN | CENPC | 3 | 738,742,744 | 1 | 0 |
| Q12824 | SNF5_HUMAN | SMARCB1 | 4 | 370,373,374,377 | 1 | 0 |
| Q16384 | SSX1_HUMAN | SSX1 | 1 | 169 | 1 | 0 |
| Q16385 | SSX2_HUMAN | SSX2 | 1 | 169 | 1 | 0 |
| Q4FZB7 | KMT5B_HUMAN | KMT5B | 2 | 352,357 | 1 | 0 |
| Q8IYW5 | RN168_HUMAN | RNF168 | 3 | 57,63,67 | 1 | 0 |
| Q8N884 | CGAS_HUMAN | CGAS | 2 | 236,255 | 1 | 0 |
| Q8NCN4 | RN169_HUMAN | RNF169 | 2 | 689,700 | 1 | 0 |
| Q96P70 | IPO9_HUMAN | IPO9 | 1 | 898 | 1 | 0 |
| Q9NQR1 | KMT5A_HUMAN | KMT5A | 2 | 229,233 | 1 | 0 |
| O60264 | SMCA5_HUMAN | SMARCA5 | 2 | 736,738 | 0 | 1 |
| Q86Y91 | KI18B_HUMAN | KIF18B | 1 | 594 | 0 | 1 |
| Q9H967 | WDR76_HUMAN | WDR76 | 2 | 215,219 | 0 | 1 |
| Q9NRZ9 | HELLS_HUMAN | HELLS | 1 | 510 | 0 | 1 |
| O15297 | PPM1D_HUMAN | PPM1D | 2 | 258,259 | 0 | 0 |
| O95760 | IL33_HUMAN | IL33 | 1 | 48 | 0 | 0 |
| P28370 | SMCA1_HUMAN | SMARCA1 | 1 | 751 | 0 | 0 |
| Q05D32 | CTSL2_HUMAN | CTDSPL2 | 1 | 283 | 0 | 0 |
| Q12830 | BPTF_HUMAN | BPTF | 1 | 1985 | 0 | 0 |
| Q149N8 | SHPRH_HUMAN | SHPRH | 1 | 796 | 0 | 0 |
| Q5H9S7 | DCA17_HUMAN | DCAF17 | 1 | 406 | 0 | 0 |
| Q69YH5 | CDCA2_HUMAN | CDCA2 | 1 | 89 | 0 | 0 |
| Q6B0I6 | KDM4D_HUMAN | KDM4D | 1 | 123 | 0 | 0 |
| Q7RTT5 | SSX7_HUMAN | SSX7 | 2 | 167,169 | 0 | 0 |
| Q7RTT6 | SSX6_HUMAN | SSX6P | 2 | 167,169 | 0 | 0 |
| Q8N531 | FBXL6_HUMAN | FBXL6 | 2 | 509,513 | 0 | 0 |
| Q8NDX5 | PHC3_HUMAN | PHC3 | 1 | 881 | 0 | 0 |
| Q8WWQ0 | PHIP_HUMAN | PHIP | 3 | 1804,1806,1808 | 0 | 0 |
| Q96B01 | R51A1_HUMAN | RAD51AP1 | 1 | 6 | 0 | 0 |
| Q96GN5 | CDA7L_HUMAN | CDCA7L | 3 | 268,273,276 | 0 | 0 |
| Q96KM6 | Z512B_HUMAN | ZNF512B | 2 | 681,684 | 0 | 0 |
| Q96ME7 | ZN512_HUMAN | ZNF512 | 1 | 340 | 0 | 0 |
| Q99909 | SSX3_HUMAN | SSX3 | 2 | 167,169 | 0 | 0 |
| Q9BVI0 | PHF20_HUMAN | PHF20 | 1 | 464 | 0 | 0 |
| Q9BXL5 | HEMGN_HUMAN | HEMGN | 2 | 35,37 | 0 | 0 |
| Q9BYG4 | PAR6G_HUMAN | PARD6G | 1 | 31 | 0 | 0 |
| Q9H1E3 | NUCKS_HUMAN | NUCKS1 | 1 | 3,6,8 | 0 | 0 |
| Q9H8M2 | BRD9_HUMAN | BRD9 | 2 | 124,127 | 0 | 0 |
| Q9NQV7 | PRDM9_HUMAN | PRDM9 | 1 | 184 | 0 | 0 |
| Q9NQW5 | PRDM7_HUMAN | PRDM7 | 1 | 184 | 0 | 0 |
| Q9NRC8 | SIR7_HUMAN | SIRT7 | 1 | 11 | 0 | 0 |

**Table 3.** Cryo-EM data collection, refinement, and validation statistics for the SHPRH-nucleosome complex.

|  |  |
| --- | --- |
|  | SHPRH-nucleosome complex (map A) |
| **Data collection and processing** |  |
| Magnification | 105,000x |
| Voltage (kV) | 300 |
| Electron exposure (e-/Å<sup>2</sup>) | 51.11 |
| Defocus range (μm) | -0.2 to -1.0 |
| Pixel size (Å) | 0.83 |
| Symmetry imposed | C1 |
| Initial particle images (no.) | 4,009,252 |
| Final particle images (no.) | 110,867 |
| Map resolution (Å) | 2.8 |
| FSC threshold | 0.143 |
| Map resolution range (Å) | 1.8-5.7 |
| **Refinement** |  |
| Initial model used (PDB code) | ModelAngelo, AF-Q149N8-F1 |
| Model resolution (Å) |  |
| FSC threshold |  |
| Model resolution range (Å) |  |
| Map sharpening *B* factor (Å<sup>2</sup>) | 58.1 |
| Model composition |  |
| Non-hydrogen atoms | 20704 |
| Protein residues | 1808 |
| Ligands | 2 |
| *B* factors (Å<sup>2</sup>) |  |
| Protein | 11.48/471.26/116.71 |
| Nucleotide | 33.89/583.81/101.82 |
| Ligand | 154.93/163.45/158.69 |
| R.m.s. deviations |  |
| Bond lengths (Å) | 0.003 |
| Bond angles (°) | 0.696 |
| Validation |  |
| MolProbity score | 1.40 |
| Clashscore | 7.26 |
| Poor rotamers (%) | 0.45 |
| Ramachandran plot |  |
| Favored (%) | 98.24 |
| Allowed (%) | 1.70 |
| Disallowed (%) | 0.06 |

### Nucleosome sliding assay

Nucleosome sliding assays were performed using 200 nM of SHPRH, 100 nM NCP with 60 base pairs of extranucleosomal DNA on one side or 30 base pairs of extranucleosomal DNA on both sides of the nucleosomal substrate, 1 mM ATP, 30 mM NaCl, 20 mM HEPES, pH 7.4 at 25°C, 0.1 mg/mL BSA, 1 mM DTT and 10% (v/v) glycerol. Reactions were incubated at 25°C and 2 µl of reaction was quenched with 10 µl quench buffer (212 ng/µL competitor DNA, 30 mM NaCl, 20 mM HEPES, pH 7.4 at 25°C, 0.1 mg/mL BSA, 1 mM DTT, and 15% (v/v) glycerol) at the indicated timepoints. 4 µL of each reaction was run on a 5% TBE gel equilibrated in 0.2x TBE running buffer and visualized by fluorescent scan at 488 nm/530 nm using a Typhoon imager (GE Healthcare). All experiments were conducted as biological triplicates.

### NADH-coupled ATP hydrolysis assay

50 µL reactions were carried out in a 364-well microplate containing 500 nM of SHPRH or *S. cerevisiae* Chd1 (residues 1-1274), 1 mM phosphoenolpyruvate (PEP), 1 mM Nicotinamide adenine dinucleotide (NADH), 250 nM NCP (30W30) or DNA (same sequence as 30W30 NCP), 5 mM ATP and 1% (v/v) pyruvate kinase/lactic dehydrogenase enzyme mix (Sigma-Aldrich P0294) in 1X buffer (50mM K·HEPES pH 7.4 at 25°C, 30 mM NaCl, 3 mM MgCl2, 1 mM DTT, and 10% (v/v) glycerol). ATP was added last, and absorbance was read at A340 nm every 10-20s on a Tecan SPARK plate reader operated at room temperature. Rates were calculated based on the slope after fitting minutes 10-20 of the sliding reactions to a linear regression. Error bars represent standard deviations. All experiments were conducted as biological duplicates.

### Figure generation, coding, and manuscript editing

Figures were generated using Adobe Illustrator, UCSF ChimeraX, and Matplotlib. Large language models were used for code generation and manuscript editing.

## Data availability

The predictome data for the H2A–H2B acidic patch is available online at https://predictomes.org/view/acidicpatch.

## Code Availability

The Python analysis code for determining acidic patch binding and all other scripts relevant to this publication are available on GitHub (https://github.com/walterlab-HMS/2024_af_h2ah2b_acidic_patch_screen).

**Extended Data Figure 1.**
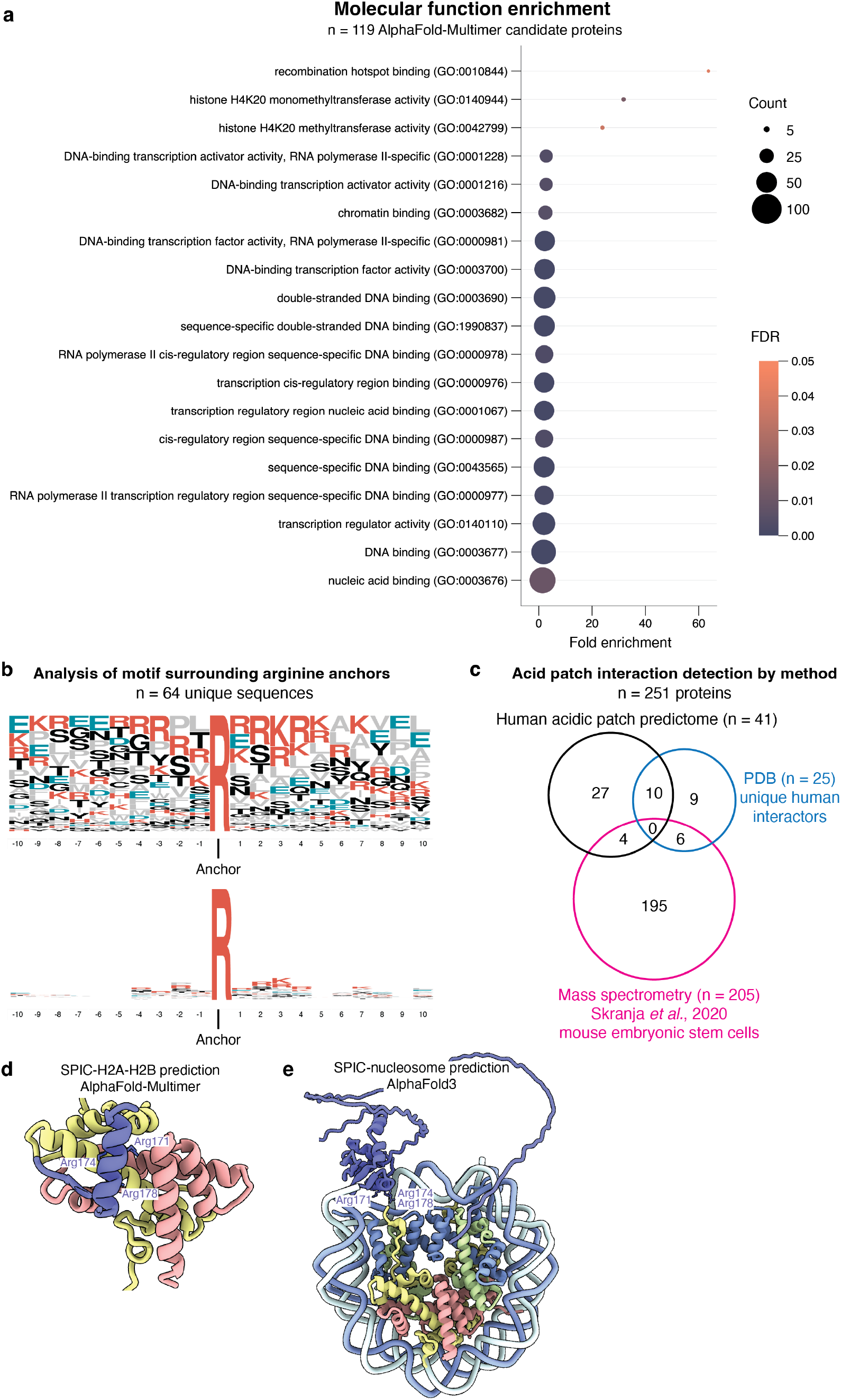
Analyzing interactors found via AlphaFold screening. **(a)** A GO based molecular function enrichment analysis of the 119 proteins that were predicted to interact with the acidic patch based on the AF2 multimer pipeline outline in Figure 2. **(b)** Sequence logo analysis of all unique sequences surrounding the identified arginine anchor residues across all 41 predictome screen hits. **(c)** Comparison of proteins that were identified as acidic patch interactors by 3 different methods (our acidic patch predictome screen), PDB structures, and mass spectrometry. **(d)** AlphaFold-Multimer SPIC-H2A-H2B prediction predicts SPIC to bind the H2A–HB acidic patch via its DNA-binding domain. **(e)** AlphaFold 3 SPIC-nucleosome prediction predicts SPIC to bind nucleosomal DNA via its DNA-binding domain.

**Extended Data Figure 2.**
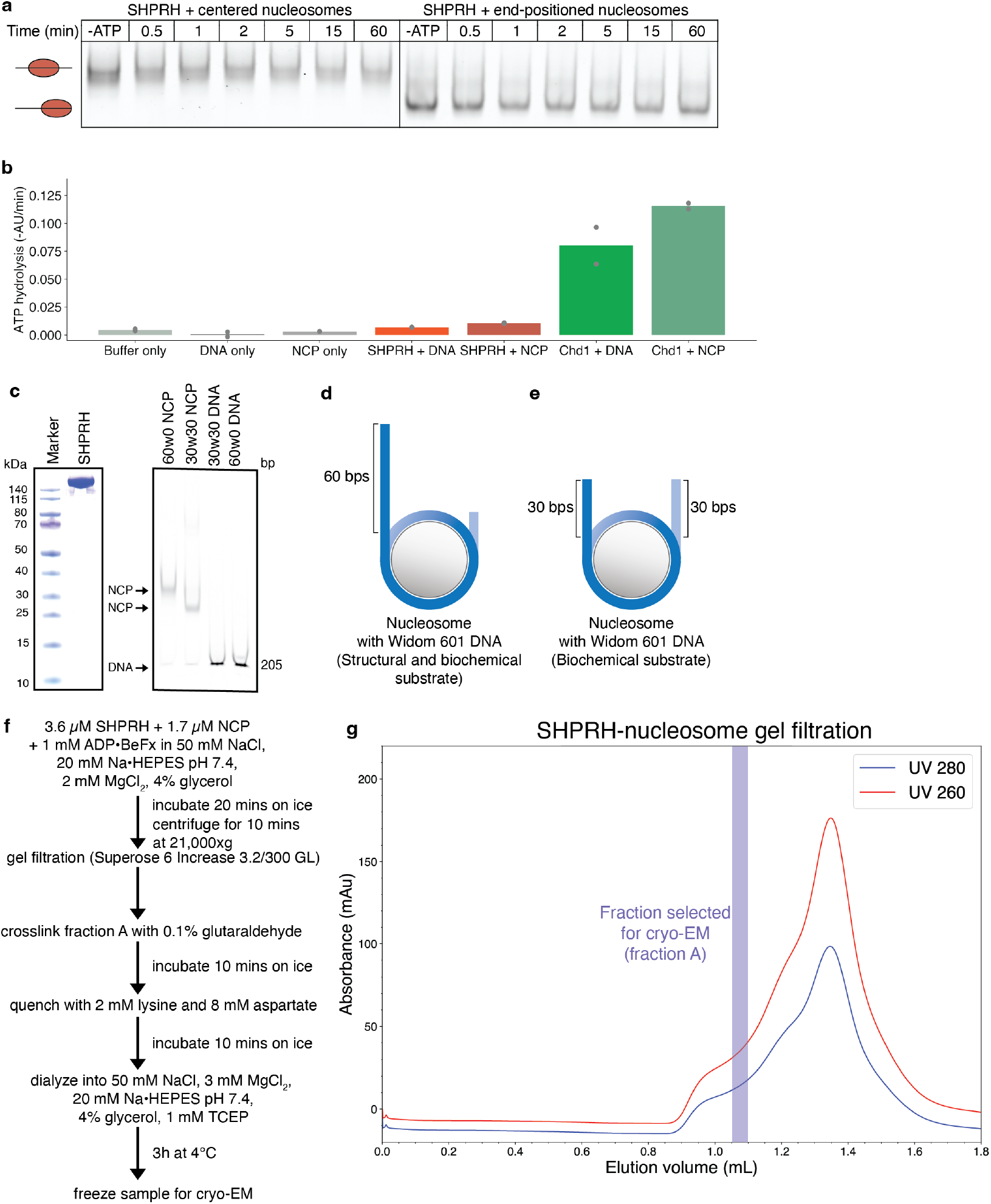
SHPRH-nucleosome complex formation. **(a)** SHPRH does not reposition end-positioned 0W60 or centered 30W30 nucleosomes. Experiments were conducted as triplicates with no clear repositioning seen in any replicate. **(b)** ATP hydrolysis rates derived from NADH-coupled ATPase assay shows DNA-and nucleosome-stimulated ATP hydrolysis activity from *S. cerevisiae* Chd1, but not SHPRH. Experiments were conducted as duplicates. Individual data points and averages are shown. **(c)** SDS-PAGE gel of purified *H. sapiens* SHPRH. The gel was stained with Coomassie Blue (OneStep Blue). **(d)** Schematic of nucleosome substrate used for structural studies, nucleosome sliding assays, and ATP hydrolysis assays (60W0). **(e)** Schematic of nucleosome substrates used for nucleosome sliding assays (30W30). **(f)** Workflow used for single-particle cryo-EM sample preparation. **(g)** Chromatogram of size exclusion chromatography run of SHPRH-nucleosome complex formation.

**Extended Data Figure 3.**
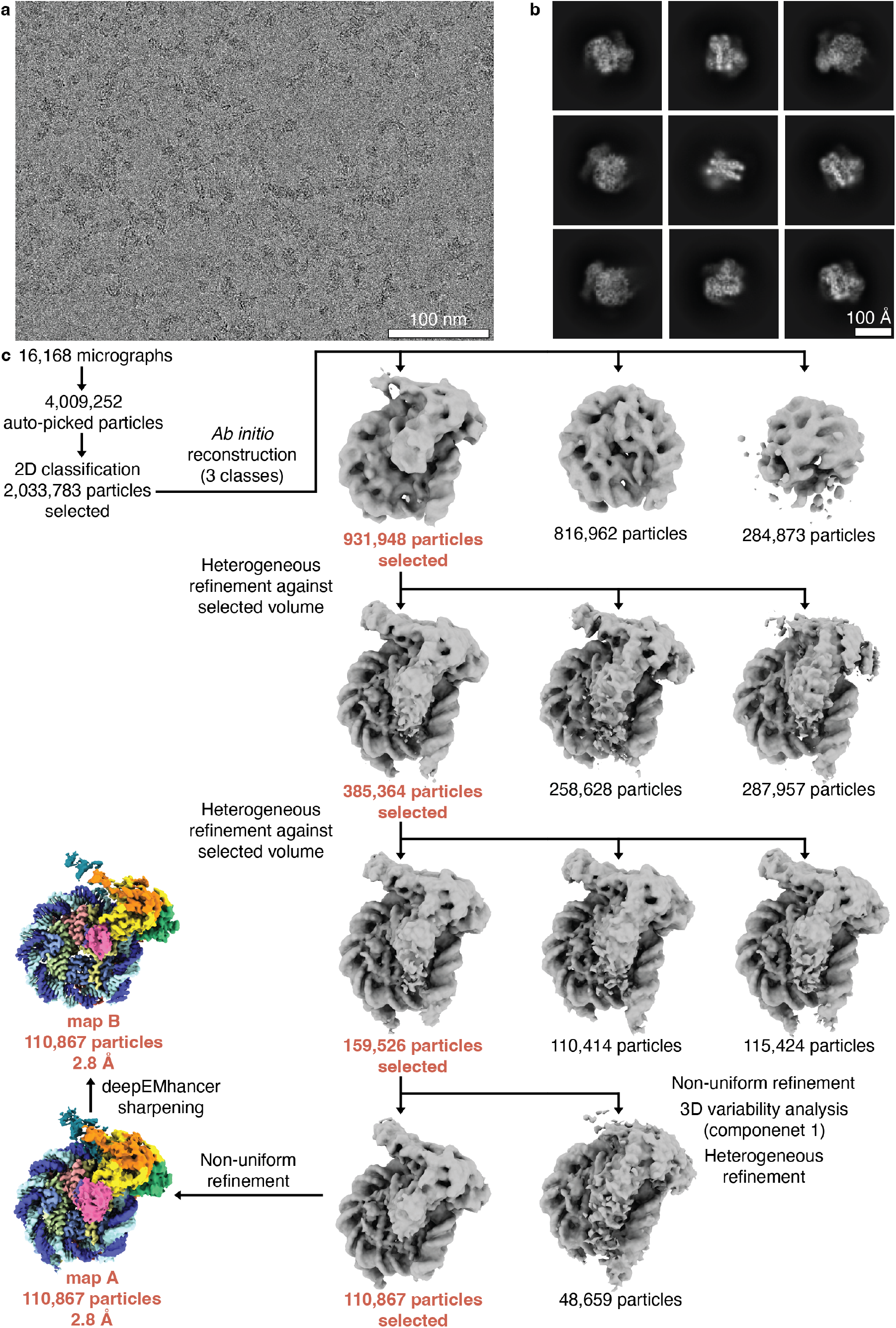
SHPRH-nucleosome cryo-EM data collection and processing. **(a)** Representative micrograph of cryo-EM data collection with scale bar (100 nm). **(b)** Representative 2D classes of SHPRH-nucleosome complexes with scale bar (100 Å). 2D classes show nucleosome- and SHPRH-like densities. **(c)** Classification tree and sorting of cryo-EM data analysis for the SHPRH-nucleosome complex. Final maps with corresponding resolutions are indicated.

**Extended Data Figure 4.**
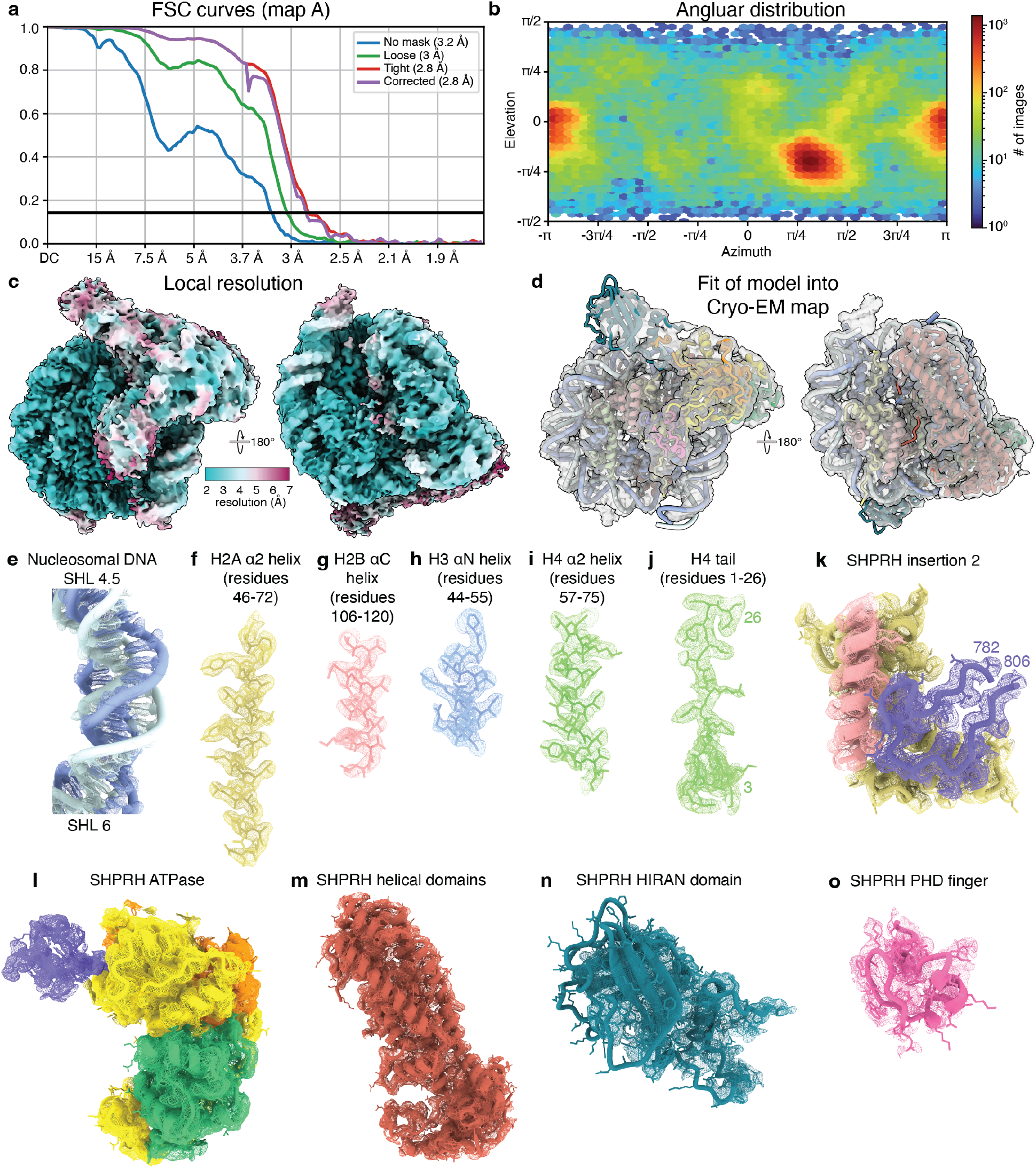
Cryo-EM map quality for SHPRH-nucleosome complex. **(a)** FSC curves of the SHPRH-nucleosome map (map A). Resolution at the FSC threshold criterion 0.143 is indicated. **(b)** Angular distribution plot of particles employed to reconstruct map A. **(c)** Local resolution of the SHPRH-nucleosome map (map A). **(d)** Fit of the SHPRH-nucleosome model into map A. **(e-o)** Cryo-EM map and corresponding model for (e) nucleosomal DNA from SHL 4.5-6, **(f-j)** indicated histone residues, **(k)** SHPRH insertion 2 (residues 782-806) with the interacting H2A-H2B dimer, **(l)** the SHPRH ATPase (resides 270-1672), (m) the SHPRH helical domains (resides 1000-1420), **(n)** the SHPRH HIRAN domain (resides 97-260), and **(o)** the SHPRH PHD finger (resides 658-709).

**Extended Data Figure 5.**
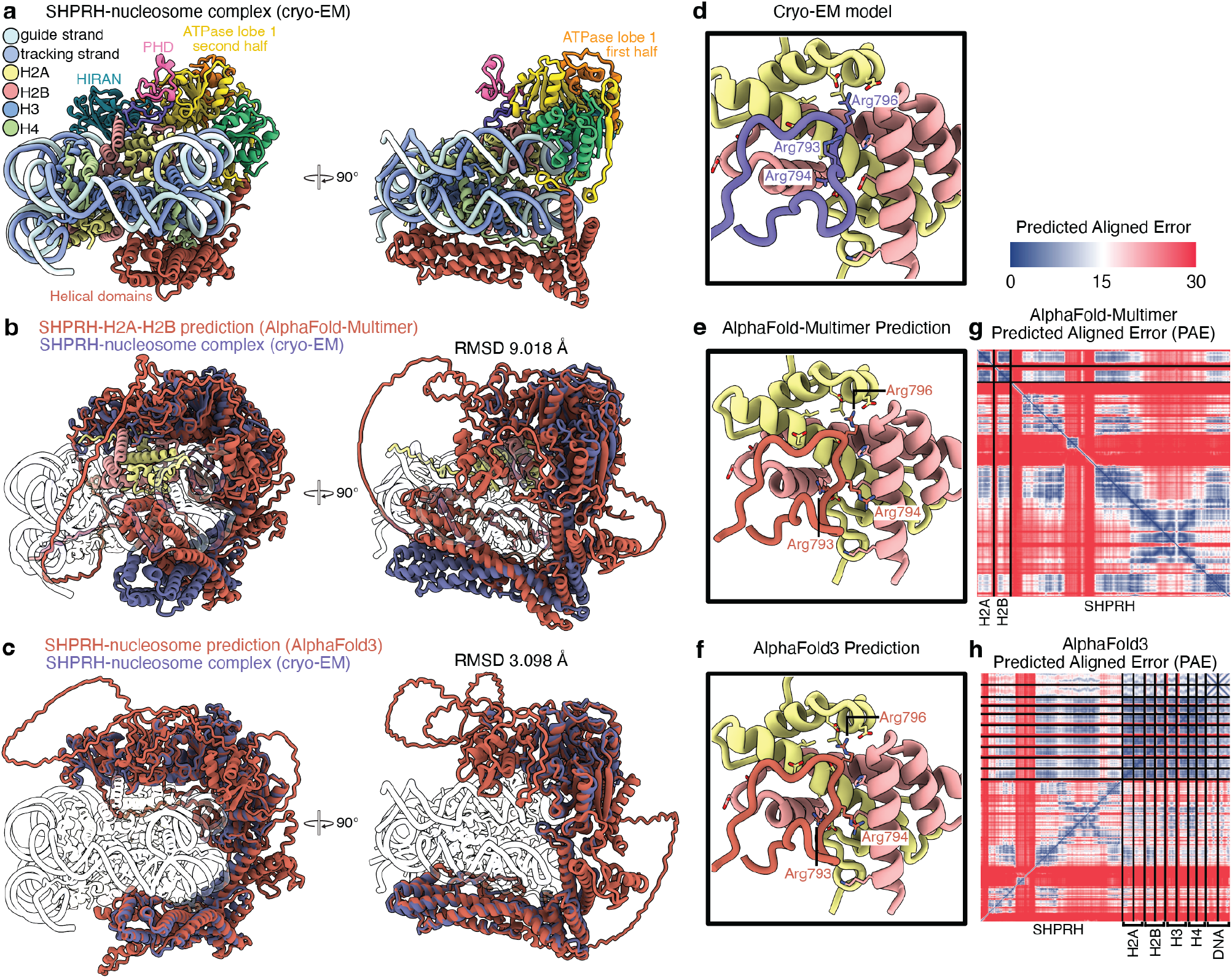
SHPRH-nucleosome cryo-EM model agrees with AlphaFold 3 prediction. **(a)** Two views of the SHPRH-nucleosome complex. **(b)** Overlay of SHPRH-nucleosome structure and AlphaFold-Multimer prediction of the H2A-H2B-SHPRH structure (model 4). Cryo-EM structure of the SHPRH-nucleosome structure is shown in purple and the AlphaFold prediction is shown in orange. **(c)** Overlay of SHPRH-nucleosome structure and AlphaFold 3 prediction of the SHPRH-nucleosome structure (model 0). Cryo-EM model of the SHPRH-nucleosome structure is shown in purple and the AlphaFold 3 prediction is shown in orange. **(d)** Positioning of SHPRH arginine anchor in our cryo-EM structure. SHPRH arginine anchor (Arg796) inserts into the nucleosome acidic patch. Arg796 and the surrounding arginines (Arg793 and Arg794) are shown as sticks. **(e)** AlphaFold-Multimer predicts SHPRH to bind the acidic patch via an arginine anchor insertion. SHPRH arginine anchor (Arg796) inserts into the nucleosome acidic patch. Arg796 and the surrounding arginines (Arg793 and Arg794) are shown as sticks. **(f)** AlphaFold 3 predicts SHPRH to bind the acidic patch via an arginine anchor insertion. SHPRH arginine anchor (Arg796) inserts into the nucleosome acidic patch. Arg796 and the surrounding arginines (Arg793 and Arg794) are shown as sticks. **(g)** Predicted aligned error (PAE) plot for model 4 of the AlphaFold-Multimer SHPRH-H2A–H2B prediction. **(h)** Predicted aligned error (PAE) plot for model 0 of the AlphaFold 3 SHPRH-nucleosome prediction.

**Extended Data Figure 6.**
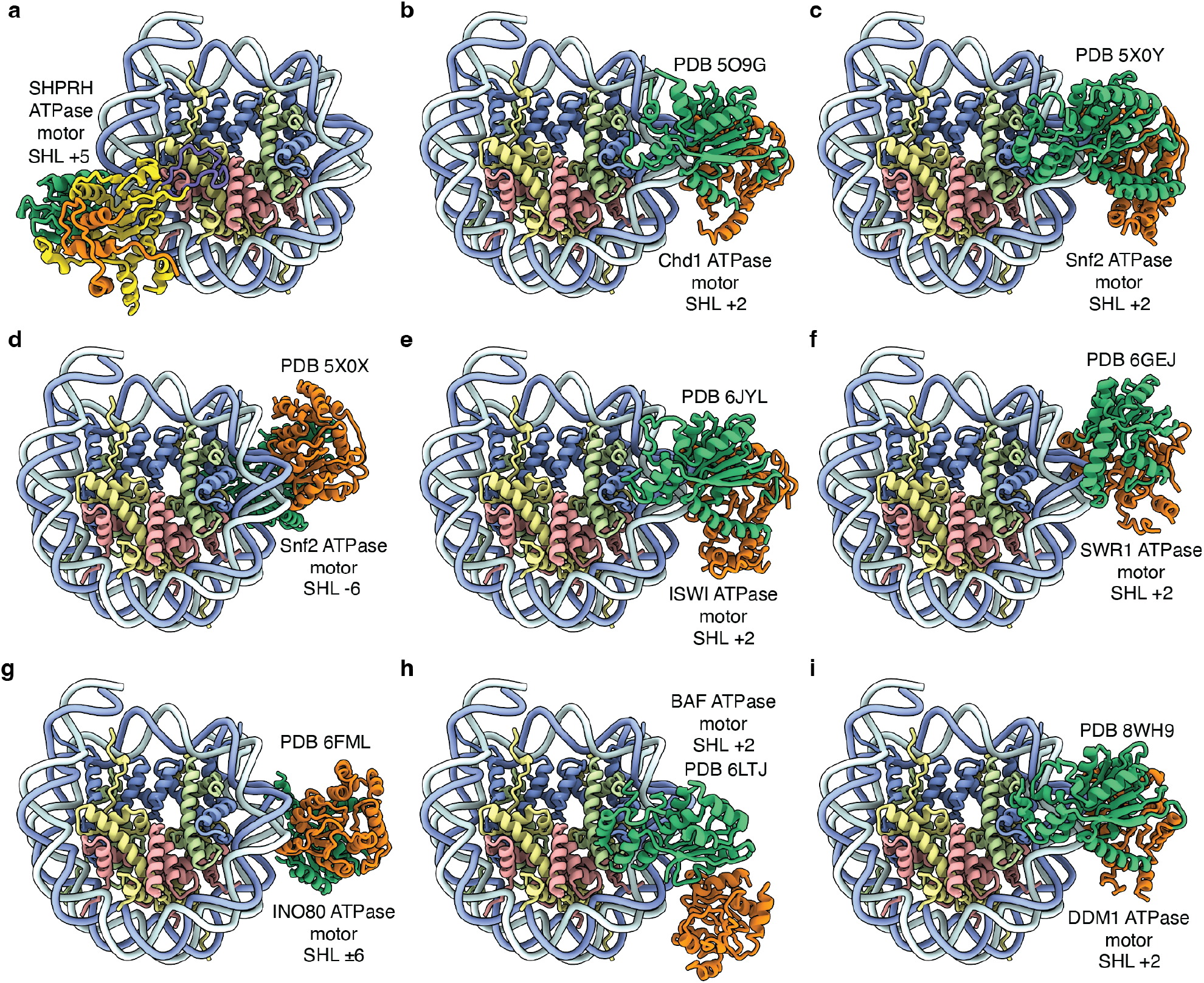
SHPRH engages the nucleosome at a different location than other SF2-type ATPases. **(a)** ATPase motor of SHPRH bound to the nucleosome. The SHPRH ATPase binds the nucleosome at SHL +5. ATPase lobe 1 is coloured in yellow and orange and ATPase lobe 2 is coloured in green. **(b)** ATPase motor of *S. cerevisiae* Chd1 bound to the nucleosome (PDB 5O9G) at SHL +2. ATPase lobe 1 is coloured in orange and ATPase lobe 2 is coloured in green. Colouring is consistent throughout. **(c)** ATPase motor of *S. cerevisiae* Snf2 bound to the nucleosome at SHL +2 (PDB 5X0Y). **(d)** ATPase motor of *S. cerevisiae* Snf2 bound to the nucleosome at SHL -6 (PDB 5X0X). **(e)** ATPase motor of the *S. cerevisiae* ISWI complex bound to the nucleosome at SHL +2 (PDB 6JYL). **(f)** ATPase motor of the *S. cerevisiae* SWR1 complex bound to the nucleosome at SHL +2 (PDB 6GEJ). **(g)** ATPase motor of the *Thermochaetoides thermophila* INO80 complex bound to the nucleosome at SHL -6 (PDB 6FML). **(h)** ATPase motor of the *H. sapiens* BAF complex bound to the nucleosome at SHL +2 (PDB 6LTJ). **(i)** ATPase motor of *Arabidopsis thaliana* DDM1 bound to the nucleosome at SHL +2 (PDB 8WH9).

**Table S1**. Table of all arginine anchors retrieved from the PDB (queried April 2024), mapped to their corresponding indices in canonical UniProt sequences as well as additional information.

**Table S2**. Table of analysis results obtained by analyzing 22,824 predicted structures representing 7,608 human nuclear proteins folded against an H2A/H2B dimer via AlphaFold multimer.

**Table S3**. Table of analysis results obtained by analyzing AlphaFold 3 predicted structures of 119 potential acidic patch interacting proteins folded against a nuclear core particle.

**Table S4**. Comparison of acidic patch interactors identified by 3 methods (predictome screen, PDB, and Mass Spectrometry) across all 7,608 human nuclear proteins run via the screen.

**Table S5.**
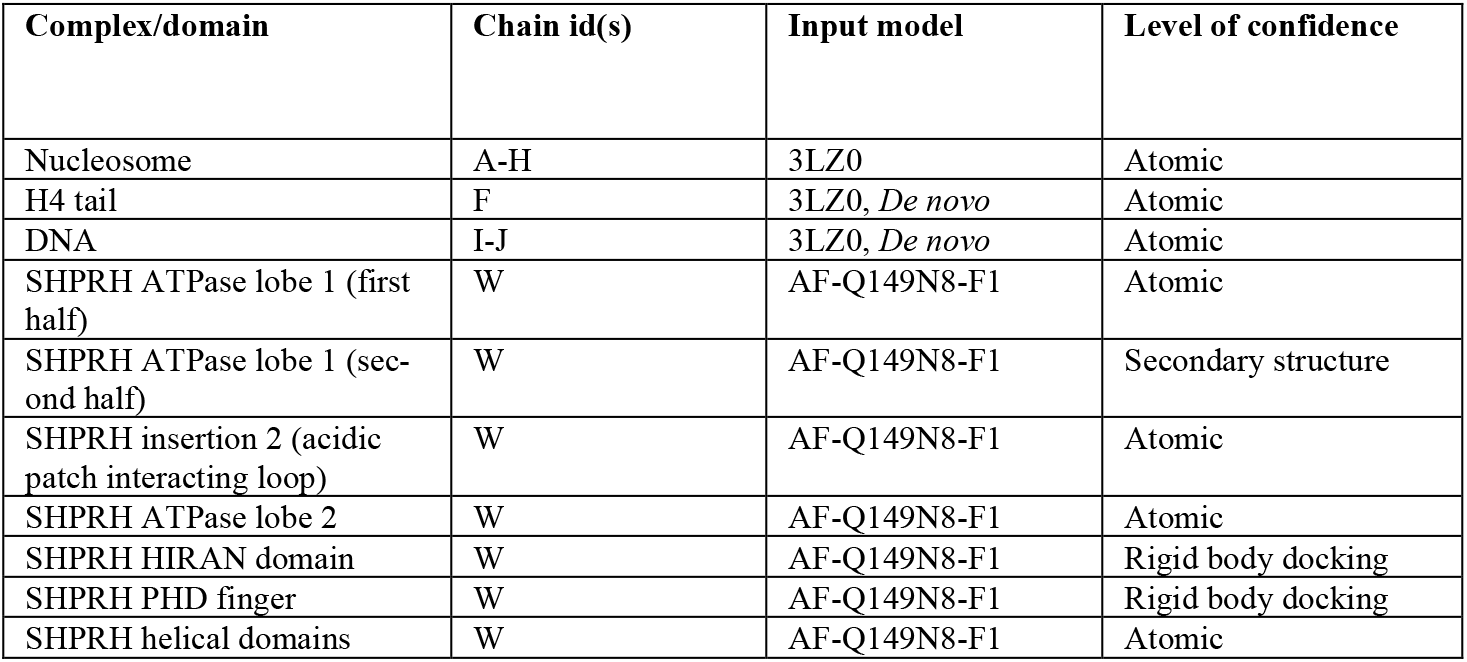
Input structural models and model confidence.

| Complex/domain | Chain id(s) | Input model | Level of confidence |
| --- | --- | --- | --- |
| Nucleosome | A-H | 3LZ0 | Atomic |
| H4 tail | F | 3LZ0, *De novo* | Atomic |
| DNA | I-J | 3LZ0, *De novo* | Atomic |
| SHPRH ATPase lobe 1 (first half) | W | AF-Q149N8-F1 | Atomic |
| SHPRH ATPase lobe 1 (second half) | W | AF-Q149N8-F1 | Secondary structure |
| SHPRH insertion 2 (acidic patch interacting loop) | W | AF-Q149N8-F1 | Atomic |
| SHPRH ATPase lobe 2 | W | AF-Q149N8-F1 | Atomic |
| SHPRH HIRAN domain | W | AF-Q149N8-F1 | Rigid body docking |
| SHPRH PHD finger | W | AF-Q149N8-F1 | Rigid body docking |
| SHPRH helical domains | W | AF-Q149N8-F1 | Atomic |

**Supplementary Video 1** | Overview of the SHPRH-nucleosome complex.

**Source Data Fig. 1** | Uncropped gels for SHPRH nucleosome sliding assays and SDS-PAGE gel of purified SHPRH.

